# Combining phase analysis and causal stimulation enables longitudinal mesoscale connectivity mapping

**DOI:** 10.64898/2026.07.09.737556

**Authors:** Leo R. Scholl, Pavithra Rajeswaran, Ryan A. Canfield, Amy L. Orsborn

## Abstract

Uncovering the circuit mechanisms of flexible behavior requires characterizing how neural signals propagate across large-scale networks and tracking how those networks evolve over time. Signal propagation through networks can be characterized by injecting activity and analyzing signals propagation, thereby measuring “causal connectivity”. However, current techniques cannot measure causal connectivity at centimeter scales over weeklong periods of time, especially in non-human primates. Here, we present a stimulation and recording method to longitudinally measure causal connectivity across centimeter-scale cortical networks in macaques. We combined multi-site optogenetic stimulation with micro-electrocorticography using a semi-chronic implant approach that enables stable daily measurements over weeks-long periods spanning years. Stimulation evoked activity reflected a mixture of local, stimulation-driven responses and secondary, delayed activity presumably driven through connections. We applied phase-based spectral analysis to disentangle stimulation-driven and secondary activity related to causal connections. We validated this approach by tracking connectivity across weeks. We demonstrated improved stability over long durations while preserving sensitivity to eyes-open vs. closed behavioral states. This platform provides a robust means to investigate the large-scale circuit dynamics underlying complex behavior in the primate brain.

## Introduction

Computations in the brain are supported by a highly recurrent network with complex anatomical organization. Understanding neural computation requires knowledge of the anatomical and functional connections that govern communication across brain regions. However, existing approaches to measure connectivity compromise between accurate estimates and scalability across brain regions. Measurements of structural connectivity map the underlying paths between neurons, but large-scale measures of structural connectivity such as tractography account for a subset of the signal flow (O’Reilly et al., 2013). In addition, structural estimates may not fully capture moment-to-moment changes in signal flow across a network because mechanisms like neuromodulation introduce dynamics across multiple timescales (Avery & Krichmar, 2017; Sweatt, 2016). Estimates of “functional connectivity” (Liu et al., 2025) more directly measure how signals move through networks by capturing how activity is shared across areas, but accurately assessing connectivity from measurements of neural activity alone remains challenging (Fig 1a). These methods are limited by their reliance on passive observations of network relationships that are inherently noisy and cannot separate causal drive from indirect correlations in a recurrent system (Das & Fiete, 2020).

**Figure 1.**
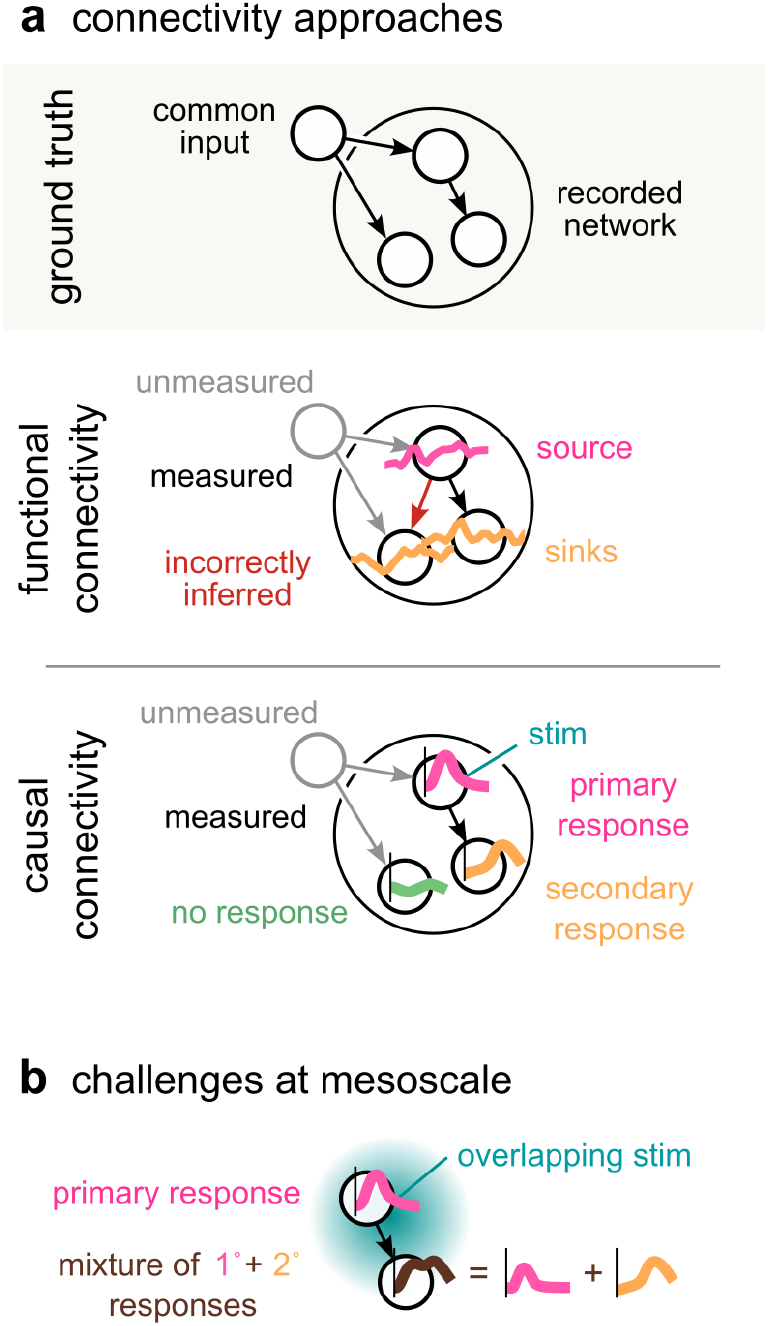
Challenges for accurately measuring network connectivity. **(a)** illustrations comparing a network (top) with connectivity inferences made using functional (middle) and causal (bottom) measurements. In functional estimates, unmeasured inputs to the network (gray) can lead to incorrectly inferred connections (red). Causal measurements experimentally introduce activity into the network via stimulation. Primary responses (pink) can be separated from secondary responses (yellow) by differences in when responses occur relative to stimulation. A site that receives no response to repeated stimulations (green) is unlikely to be connected to the stimulation site. **(b)** Mesoscale signals that combine the activity of many neurons may contain mixtures of primary and secondary responses (brown), increasing the difficulty of spatially and temporally resolving connectivity.

The flow of signals through a network is thought to be more accurately assessed using stimulation, which directly measures if an evoked signal at one site is received at another (Bauer et al., 2018; Bundy et al., 2023). The resulting estimates are sometimes called “causal connectivity” due to the external, causal experimental manipulation. Causal connectivity has successfully identified microscale circuitry at the level of single neurons (Finkelstein et al., 2025; Guo et al., 2009; Histed et al., 2009; Randi et al., 2023; Russell et al., 2022) and specific long range connections (Bundy et al., 2023; Qiao et al., 2020; Yazdan-Shahmorad et al., 2018). Scaling these methods to measure entire networks, however, has proved challenging, especially in larger brains. Widespread optical recording and stimulation of neurons requires genetic engineering methods that aren’t widely available for large animals like primates (Feng et al., 2020; Izpisua Belmonte et al., 2015), limiting their application and translation to humans. Electrical stimulation creates undesirable recording artifacts when paired with electrophysiology and indiscriminately drives axons (Balbinot et al., 2025; Histed et al., 2009; Nowak & Bullier, 1998), complicating accurate estimates of connectivity.

Combining stimulation and recording techniques presents further challenges for resolving the temporal dynamics of connectivity. Whole-brain functional imaging such as fMRI doesn’t have the temporal resolution to identify sub-second changes within a brain area (Glover, 2011; Oya et al., 2017; Petkov et al., 2015), especially when combined with compatible stimulation methods (Valero-Cabré et al., 2005). Further, measurements over long timescales are needed to detect changes in connectivity related to learning, for instance, but few methodologies provide the ability to record and stimulate the same neurons across long periods. Repeatedly positioning electrodes for each measurement can be time consuming and imprecise, and existing chronic approaches suffer from photoelectric or axonal stimulation artifacts (Cho et al., 2022; Hong et al., 2025) or don’t yet scale past millimeter coverage (Finkelstein et al., 2025).

Finally, the analytical approaches used to compute causal connectivity pose challenges to making large-scale measurements. Existing methods to measure causal connectivity often use latency to distinguish between direct, stimulation evoked activity and secondary, network driven responses (e.g., Banerjee et al., 2010). This separation has been successful for spatially local neural measurements (e.g., single neurons), which generate completely independent signals. Mesoscale signal modalities, which combine the activity of thousands of neurons (e.g. local fields), may be advantageous for detecting sparse connections across large networks because they densely sample neural populations (Buzsáki et al., 2012). But these signals are less spatially independent; stimulation-evoked activity may overlap with secondary responses, making estimates like latency impractical (Fig 1b). Approaches that can separate stimulation-evoked and secondary responses at the network level will enable measurements across a larger class of recording modalities.

Here we address this combination of methodological and analytical challenges to advance mesoscale measures of causal connectivity. First, we developed a technique in macaques to repeatably apply optogenetic stimulation across multiple cortical regions (centimeter-scale) combined with micro-electrocorticography (µECoG) recordings. µECoG can record activity with high spatial and temporal resolution and with large spatial coverage (Ouchi et al., 2025; Trumpis et al., 2021). We combined µECoG recordings with optogenetic stimulation to avoid electrical stimulation artifacts and directly target excitatory cells without stimulating passing axons. Second, we applied analytical methods based on phase differences to resolve the mixture of stimulation-evoked and connectivity-driven responses inherent to mesoscale signals like µECoG. We present the causal connectivity measured across motor cortices in two macaque monkeys and describe the advantages of our method over functional connectivity estimated from resting data. We show that causal connectivity reveals a distinct network of connections compared to functional measures. Using our combined stimulation and recording approach, we further demonstrate the ability to maintain consistent measurements of connectivity across days to weeks yet detect meaningful differences in connectivity between two simple behavioral states.

## Methods

### Surgical procedures

All procedures were approved by the Institutional Animal Care and Use Committee at the University of Washington. Two male rhesus macaques (Monkey 1: B, 9 years old; and Monkey 2: A, 10 years old) were implanted with multi-modal recording chambers as described previously (Ouchi et al., 2025). The chambers were designed to precisely locate a molded silicone artificial dura spanning portions of primary motor cortex (M1), premotor cortex (PM), and prefrontal cortex (PFC). Headposts were attached with bone screws posterior to the recording chambers to enable head fixation and serve as electrical references.

The craniotomy and durotomy were made while the animals were under propofol anesthesia. Following durotomy, adeno-associated virus targeting excitatory cells was injected at sites across the cranial window to broadly express channelrhodopsin and a fluorescent reporter (AAV9-CaMKIIa-hChR2(H134R)-mCherry gifted from Karl Deisseroth; Addgene viral prep #26975-AAV9). An artificial dura with 32 holes arranged in a grid pattern was inserted into the chamber to locate injection sites. A Hamilton syringe (#7105) mounted in a microinjection syringe holder (model 5000, Kopf Instruments) was angled perpendicular to the cortical surface at the center of the chamber. Injections were made at 7 (Monkey 1) or 8 (Monkey 2) sites, at depths of 3, 2, and 1 mm from the cortical surface. Sites were chosen to maximize the spatial extents of viral expression while avoiding major blood vessels. For each injection, the syringe was depressed manually over 30 seconds to inject 1.0 µl of AAV vector. Following each injection, the syringe was kept in place for one minute before advancing to the next depth, or ten minutes following the final depth at each site. Following injections, the chambers were hermetically sealed using screws and water-tight sealing gaskets (Orsborn et al., 2015) and maintained with regular cleanings. Animals were trained using positive reinforcement to direct load into a primate chair fitted with an adjustable head restraint (Hybex Innovations, Montreal QC). Under sterile conditions, the recording chamber was opened temporarily for inspection, imaging, or for semi-chronic implanting of recording electrodes and fiberoptics.

### Viral expression assays

Expression of AAV was verified with *in vivo* fluorescence imaging (Monkey 1, 1 year post-injection; Monkey 2, 1 month post-injection). A filter cube (Thorlabs) was attached to the primate chair using a Magic Arm (Manfrotto 244). A multi-wavelength filter set (Brightline) was used in combination with a cold white LED array (Thorlabs LIUCWHA) and a 25 mm lens (Arducam LN041) both mounted to the exterior using 3D printed fixtures. We used a Raspberry Pi HQ camera (Raspberry Pi Foundation) to capture images. The Python package pi-macroscope (github.com/leoscholl/pi-macroscope) was used to trigger a 30-second exposure with the room lights turned off, revealing mCherry fluorescence. Only the red channel of each image was used to measure expression.

One animal (Monkey 1) was euthanized at the end of study to verify the extent and cell-type specificity of viral expression in the chamber. Cardiac perfusion was performed with normal saline followed by a 4% solution of paraformaldehyde in phosphate buffered saline. The brain was extracted and postfixed for 24 hours before being cryoprotected in a 30% sucrose solution. A block of tissue around the chamber was sliced on a freezing microtome at 50 µm thickness. Fluorescence signals for mCherry (primary antibody 1:250, mouse anti-mCherry, Takara 632543; secondary antibody 1:500, Invitrogen) and parvalbumin (PV, primary antibody 1:10, rabbit anti-PV, Swant PV27, secondary antibody 1:500, Invitrogen) were detected immunocytochemically at four sections across two injection sites. At one injection site, an additional two 30 µm sections were collected for in-situ RNA hybridization (RNAscope Multiplex Fluorescent Detection Kit v2, ACD 323110) to detect excitatory cells expressing vesicular glutamate transporter 1 (VGlut, ACD 492791). For pretreatment, sections were dried, washed, and dehydrated on Superfrost slides (Fisher 1255015) followed by heat-induced target retrieval and protease III digestion. Both sections were hybridized with VGlut (RNAscope probe SLC17A7, opal 1:750, Akoya FP1487001KT). One section was co-labeled for mCherry (primary antibody 1:250, mouse anti-mCherry, Takara 632543; secondary antibody 1:500, Invitrogen) and the other was co-labeled for RFP (primary antibody 1:250, rabbit anti-RFP, Rockland 600-401-379; secondary antibody 1:500, Invitrogen). All sections were counterstained for DAPI (1:5000, Thermo Fisher D1306) and coverslipped with polyvinyl alcohol mounting medium with DABCO (Sigma Aldrich 10981). Images of PV-stained sections were acquired on an epifluorescence microscope at 20x magnification (Leica DM5500B). Images of VGlut-stained sections were acquired on a laser-scanning confocal microscope (Leica SP8X). Cell counting was performed manually using QuPath.

### Simultaneous electrophysiological recordings and optogenetic stimulation

Neural signals were recorded using a 244-electrode µECoG array (762 µm inter-electrode pitch and 229 µm contact size) embedded in a silicone artificial dura to enable registration within the implanted chamber (Chiang et al., 2020; Orsborn et al., 2015; Ouchi et al., 2025). µECoG signals from 240 electrodes were amplified and digitized at 25 kHz using an eCube recording system (White-Matter Inc., Seattle WA).

The µECoG array was designed to be optically opaque but with a grid of 32 holes that allow access to sites across the cortical surface. This hole pattern matched the artificial dura used during viral injection surgery. Fiberoptic cannula (200µm diameter, 0.37NA; Hangzhou Braintech Technology Co., Xi’an City) were aligned to these holes using a custom fixture secured atop the µECoG array with silicone adhesive. The optical path from each fiber cannula to the underside of the array was verified under a stereo microscope with illumination from below prior to sterilization in hydrogen peroxide vapor (STERRAD) and implantation.

At the start of each insertion period (see Data selection), the combined, sterilized µECoG array and fiberoptic fixture were inserted into the recording chamber during a sterile procedure performed while the animal was awake. The µECoG assembly included protrusions which were registered to matching notches in the chamber to provide consistent positioning and orientation (Orsborn et al., 2015). On each recording day, animals were seated with head restraint in a dimly lit recording booth. One of the 32 fiberoptic cannula (a “stimulation site”) was then connected to a 450 nm (blue) laser coupled to a 200 µm diameter optic fiber. The laser was developed in-house by the Instrumentation Services Core at the Washington National Biomedical Research Center. Stimulation timing was controlled by a digital signal which was recorded by the eCube recording system along with the neural data to facilitate synchronization. Laser power was set to achieve 10 ± 2 mW at the fiber tip measured prior to sterilization using a calibrated photodiode sensor (Thorlabs S121C).

Stimulation was repeated with Poisson-randomized inter-stimulation intervals (ISI) and random pulse width drawn uniformly from a set of pulse widths (Monkey 1: [1, 3, 10, 15, 20] ms; Monkey 2: [3, 10, 30] ms) to avoid habituation and provide a broad-spectrum network drive (rather than a singular frequency) (Qiao et al., 2020; Yang et al., 2018, 2021). ISI were drawn from an exponential distribution with mean 500 ms, then any ISI below 100 ms was discarded to avoid overlapping events.

### Data selection

Our implants enabled semi-chronic insertions of µECoG and fiber optics lasting from 2 – 21 days. Insertions began based on assessment of tissue inside the recording chamber and ended based on estimates of tissue health following prior insertions and on assessments of signal quality. Recordings were made on all days when the µECoG array and fiberoptic assembly were implanted. On a given recording day (*N* = 63 days for Monkey 1 and 220 days for Monkey 2), a subset of the 32 stimulation sites were chosen for data collection (Monkey 1 mean 4.3, range 1 - 9 sites per day; Monkey 2 mean 2.8, range 1 - 23 sites per day). We recorded 350 and 600 stimulation trials per site each day for Monkey 1 and Monkey 2, respectively. On at least one occasion for each animal, data for each of the 32 stimulation sites were collected within a single insertion (spread across 14 days in Monkey 1; 2 days in Monkey 2). Data within this single insertion was used for across-site comparisons for each monkey. For longitudinal analyses, we split data into multi-day series during which the same stimulation site was targeted repeatedly. Data were considered to belong to the same series if there were no more than four days between recordings. This approach combined data across insertions in some cases (6 / 20 series spanned multiple insertions for Monkey 1; 7 / 55 series for Monkey 2).

### Evoked µECoG responses to laser stimulation

For each data collection session, each channel was mean subtracted, low-pass filtered beneath 500 Hz with a fourth-order Butterworth filter and decimated to 1 kHz sampling rate. Evoked activity in the 250 ms preceding and following each laser pulse were identified using the digitized photodiode signals from the laser. Evoked activity was normalized by subtracting the mean activity in the 250 ms preceding laser onset, forming a (time, electrodes, trials) tensor. Outlier trials were removed based on root mean square (RMS) voltage in the 250 ms preceding laser onset. Trials during which more than 10% of electrodes exceeded 5 SD from the median RMS were excluded from further analysis.

We estimated the single-trial response to stimulation for each electrode as the maximum evoked activity (positive or negative) over the 250 ms following laser onset, resulting in a (electrodes, trials) response matrix. We visualized the spatial map of responses to stimulation using the mean of the response matrix across time. We quantified the total size of the evoked response across the µECoG array by calculating the mean response across electrodes. We also generated a chance distribution using 30 random samples of time-series data from non-stimulated epochs, each with the same number of trials as the stimulation data. A one-tailed Wilcoxon signed rank test was used to compare the observed mean response to the null distribution for each stimulation site, corrected for multiple comparisons using false discovery rate correction with α=0.05. To aide visualization, spatial data at each electrode was scaled by the p-value from this comparison (Ouchi et al., 2025).

### Spatial correlation analysis

We analyzed the consistency between spatial maps by calculating the pixelwise correlation between pairs of maps. Data maps were normalized by their magnitude prior to computing correlation to control for variability in overall signal amplitude while preserving spatial relationships across electrodes. All correlation coefficients were calculated using Pearson’s method.

### Latency-based connectivity estimation

We analyzed connectivity using latency-based metrics that estimate the timing delay between stimulation and neural responses. Evoked responses were first band-pass filtered using a complex-valued multi-taper filter to isolate the high gamma (50 – 200 Hz) band (three Slepian tapers, length 30 ms with 150 Hz bandwidth and 125 Hz center frequency). Power was computed as the magnitude of the signal after convolving the evoked potentials with the filter, preserving the original shape of the data and centering the output.

Stimulation response latency was computed from the high gamma band power using an established signal detection method (Qiao et al., 2020). This approach computes a time-varying likelihood of whether data belongs to one of two conditions (null or stimulation) and uses a signal detection theory approach to determine the latency at which the stimulation condition appears in the data. Importantly, this approach controls for different response magnitudes when computing latency (Banerjee et al., 2010). The evoked (time, electrodes, trials) tensor was split into null (the 250 ms preceding stimulation) and stimulation (the 250 ms following stimulation) conditions. The log-likelihood ratio (LLR) for each timepoint was computed as

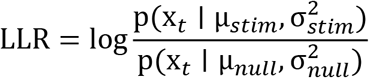

where x*_t_* is the data (from null or stimulation condition) at time t for a given trial, μ*_stim_* and *σ^2^_stim_* are the mean and variance, respectively, of all stimulation condition data across all other trials, and μ*_null_* and *σ^2^_null_* are the mean and variance, respectively, of null condition data across all other trials. A cumulative sum on the LLR timeseries produced the accumulated log-likelihood ratio (AccLLR):

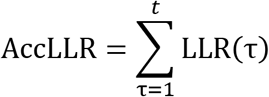

The threshold for detecting stimulation activity was selected using receiver operating characteristic (ROC) analysis. The probability of correct detection was calculated at each threshold *t*ℎ*r* as the proportion of trials on which either AccLLR_stim_ exceeded +*t*ℎ*r* or AccLLR_null_ fell below −*t*ℎ*r*. Similarly, the probability of incorrect detection counted trials when AccLLR_stim_ fell below −*t*ℎ*r* or AccLLR_null_ exceeded +*t*ℎ*r*. Finally, the threshold was chosen that maximized the difference between correct and incorrect detection of stimulation across trials. Latency for each trial was computed as the time at which the threshold was exceeded on each trial. Area under the curve (AUC) at the best threshold was computed using the DeLong method (Sun & Xu, 2014) and a *p*-value for each channel was computed under the null hypothesis that the AUC = 0.5 using a one-sided Z-test. Following false discovery rate correction with α=0.05, only electrodes with AUC significantly above 0.5 were included as having a statistically significant latency.

To control for the response magnitude influencing the time at which it is detected, we gradually introduced gaussian noise until all significant responses had a similar AUC, meaning they were equally well detected. First, out of all electrodes with a significant latency, we identified the electrode with the smallest AUC. For each remaining electrode with a significant latency, the latency analysis was repeated iteratively, but with additional noise added in steps of 0.1 SD. Whenever an electrode had AUC at least as small as the minimum AUC, it was removed from further iterations.

We used latency estimates to compute connectivity using the stimulation evoked response ratio (SERR; Yazdan-Shahmorad et al., 2018). First, we manually selected a cutoff to exclude responses below a certain latency. The cutoff was chosen such that the majority of responses at the stimulation site, presumed to include direct responses, were excluded. SERR was calculated by dividing the evoked response at a given electrode by the mean evoked response at the four electrodes closest to the stimulation site. SERR was set to zero at electrodes where the computed latency was less than the cutoff. SERR was also set to zero at electrodes where ROC analyses did not detect a significant response.

### Stimulation-locked imaginary coherence

We estimated connectivity using statistical relationships between pairs of electrodes using multiple metrics: imaginary coherence and Grainger prediction (see below). Imaginary coherence is similar to the magnitude of coherence but with the convenient property that signals with zero phase difference are ignored (Nolte et al., 2004). Because the stimulation sites were not positioned directly above a single electrode, we computed the imaginary coherence between the four electrodes nearest to the stimulation site (considered *stimulation electrodes*) and other electrodes in the array that were not directly adjacent to the stimulation. We averaged the four values to get an estimate of the imaginary coherence to the stimulation site. Two Slepian tapers of length 60 ms with a bandwidth of 25 Hz were used to estimate spectral coefficients. Imaginary coherence (IC) was computed for a given pair of signals 1 and 2 with

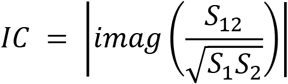

where *S*_1_ and *S*_2_ are the two analytical signals and *S*_12_ is their cross spectrum, computed as means over trials and tapers. The phase difference between the two signals was computed as the angle of the complex valued *S*_12_.

We analyzed two time windows to identify the change in IC induced by stimulation. We call this difference the stimulation-locked imaginary coherence (SLIC):

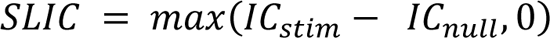

where *IC_stim_* is the imaginary coherence of the spectral estimate in the 60 ms after stimulation and *IC_null_* is the imaginary coherence of the spectral estimate of the 60 ms before stimulation. Thus, SLIC measures the increase in imaginary coherence following stimulation. To test for significance, we generated a null distribution of SLIC values for each stimulation site using data recorded during non-stimulated epochs and selecting time points randomly. A one-tailed Wilcoxon signed rank test was used to compare the observed SLIC at each electrode to its corresponding null distribution, corrected for multiple comparisons using false discovery rate correction with α = 0.05.

### Granger prediction

We used spectral granger prediction (GP; Geweke, 1982) to analyze connectivity in the absence of stimulation using two different datasets: without stimulation while the animal was at rest, and during stimulation applied to control sites (lacking AAV expression). For analyses without stimulation (GP), data were aligned to random timepoints, during which no stimulation occurred, spaced in time using the same Poisson distribution as the stimulation pulse trains (mean time between events 500 ms, with a refractory period of 100 ms). Fourier coefficients were computed using the multi-taper method with the same parameters as used to calculate SLIC. We then used the Spectral Connectivity python package (Denovellis et al., 2022) to compute pairwise spectral granger prediction from each of the four electrodes surrounding a stimulation site to all other electrodes, averaging the result from each of the four source electrodes. To test for significance, we compared actual Granger values to a chance distribution using a one-tailed Wilcoxon signed rank with false discovery rate correction with α = 0.05 to correct for multiple comparisons across electrodes. Each chance distribution contained 30 values of GP generated by scrambling the time axis repeatedly, destroying the temporal relationships across signals (Cohen, 2014).

We also computed stimulation-evoked granger prediction (SEGP) to test the importance of the specific analysis being used. We followed the same procedure as GP on but replaced the (time, electrodes, trials) rest data tensor with the 60 ms following laser stimulation.

### Trial adding analysis

We estimated the number of trials necessary to obtain a consistent stimulation-driven response using a trial adding analysis. For each stimulation site across each insertion period, we concatenated all the trials of stimulation data. We focused this analysis on subsets of our data (sites and insertions) with at least 3,375 trials, which ensured sufficient data to divide into at least 5 groups of at most 675 trials. The data was divided into increasing number of trials without replacement, in increments of 25 trials, and we computed the evoked response map for each dataset (batches of 675 trials) as we varied the number of trials used. We estimated map consistency by calculating the correlation coefficients across all possible pairs of maps for each data batch with the same trial size. Lastly, we calculated the maximum of the mean correlation at each trial size, then found the number of trials it took to reach 90% of that maximum.

### Connectivity metric comparisons and assessments

We used a resampling approach to estimate the similarity of maps generated within and across different metrics of connectivity. For each stimulation site with significant evoked responses to laser stimulation, we randomly drew half the trials without replacement to create two subsets of the data. We then calculated maps of connectivity for each data subset using SLIC and SEGP. We followed the same procedure to calculate two maps of connectivity at the same stimulation site using GP, substituting data where stimulation was applied to control sites with no significant evoked response. We computed the spatial correlation between the two maps generated by each method as a measure of within-metric consistency. We also computed the spatial correlation across maps generated by different metrics as a measure of across-metric consistency. Finally, we repeated the entire procedure 30 times using different random seeds each time to generate a distribution of within- and across-metric comparisons.

We also used this resampling approach to estimate the consistency of each connectivity metric across frequency bands. We computed connectivity in five frequency bands for each of the two maps generated in each resample in the previous analysis. We then calculated the spatial correlation within and across different frequency bands for each connectivity metric.

### Spatial distribution of connectivity

We analyzed the relationship between connectivity and distance to compare the spatial relationships uncovered by SLIC and GP connectivity metrics. For each stimulation site and each frequency band individually, we constructed a histogram of the significant connections identified from that site, dividing the data into 20 bins by the distance between electrode pairs. We normalized the cumulative distribution of each histogram by subtracting its minimum connectivity and dividing by its sum, generating a metric that captures “relative connection strength”. We then compared the distributions of short-and long-range connections. The minimum and maximum distance between electrode pairs in our data was 0.8 mm and 12 mm, respectively. We counted each individual connection strength as short-range if the distance between the stimulation site and the recording electrode was less than 4 mm and long-range if the distance was greater than 8 mm.

We also directly compared the distributions of connectivity over distance between SLIC and GP to non-parametrically test whether the metrics were different for each frequency band. We pooled data across all sites with significant stimulation-evoked responses. Then we constructed a histogram of connection strength, binned by distance, for each frequency band using SLIC and GP. We compared the cumulative distributions of each metric using a Kolmogorov-Smirnov (KS) test.

### Longitudinal stability

We evaluated the stability of optogenetic response, SLIC connectivity, and GP connectivity using a pixelwise Pearson correlation between two maps measured at different time-points. We then compared the distributions of correlation coefficients on days of stimulation to chance distributions using Mann-Whitney U tests. For stimulation data, the chance distributions were drawn from resting data with no stimulation. For GP analysis, we use chance distributions created by shuffling the time axis repeatedly for each epoch.

### Eye behavior analysis

The animals’ eye positions were monitored and recorded during all data collection sessions. Images of both eyes were captured using an infrared camera at 240 Hz (FLIR CM3-U3-13Y3M-CS). Pupil positions were estimated online using Oculomatic software (Zimmermann et al., 2016) and recorded as analog voltages to the eCube recording system.

Animals sometimes closed their eyes for minutes at a time during laser stimulation epochs. We estimated the times at which the animal had both eyes open or closed using eye tracking data by analyzing the 500 ms preceding and 500 ms following each laser onset. We considered the eyes closed if eye position remained unchanged for more than 80% of that time, and open if eye position remained unchanged for less than 50% of that time. For each eye, we considered position unchanged if it remained within a range of three pixels for at least 10 ms.

We used this parcellation of data to analyze whether the connectivity we measured using stimulation was affected by the animals’ behavioral state. We estimated bootstrapped distributions of connectivity during eyes open and eyes closed epochs using a resampling procedure. We focused these analyses on the subset of our data with longitudinal measurements where five stimulation sites were repeatedly stimulated for more than 5,000 trials. We randomly selected groups of 200 trials without replacement, separately for trials on which the animals’ eyes were open or closed. We calculated a SLIC connectivity map in the 12-150 Hz band for each group of trials and repeated this process 50 times to create eyes open and eyes closed distributions of connectivity. We compared these empirical distributions using d-prime, a measurement of discriminability between two distributions (Green & Swets, 1988; Rouse et al., 2013). Finally, we repeated the entire process 50 times with shuffled eye labels to create a null distribution of d-prime values. We compared observed values of d-prime to the null distribution using a one-tailed Wilcoxon signed rank test, corrected for multiple comparisons across electrodes using false discovery rate correction. To aide visualization, the opacity of the connectivity difference at each electrode was scaled by the observed d-prime at that electrode:

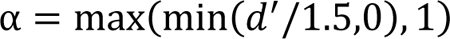

such that full opacity indicates a d’ greater or equal than 1.5 and full transparency indicates a d’ of zero.

## Results

We designed a semi-chronic approach to combine micro-electrocorticography (µECoG) with fiberoptics for repeatable optogenetic stimulation across days in monkeys. We recorded neural activity using a 244 electrode µECoG array and stimulated using a fiber-coupled laser at 32 sites interspersed within the 12 x 12 mm array (Fig 2a). We made viral injections of AAV to express channelrhodopsin (ChR2) and a fluorescent reporter across motor cortices in two monkeys (Fig 2b). Fluorescence imaging revealed a pattern of expression that was visually consistent with the injection locations. ChR2 expression was targeted to excitatory cells using a CamKII promoter (Watakabe et al., 2015). In one animal we tested whether infected cells were excitatory or inhibitory at two injection sites using in-situ labeling (Fig 2c). We verified that most identified infected cells were glutamatergic (274/308 mCherry+ cells were VGlut+ summed across two sections, 95% confidence interval: 85.46% to 92.46% excitatory) and that most identified infected cells were not parvalbumin expressing (PV) inhibitory cells, one of the most common inhibitory neuron subtypes (32/834 mCherry+ cells coexpressed PV+ summed across four sections, 95% confidence interval: 2.53% to 5.14% PV; Fig 2d).

**Figure 2.**
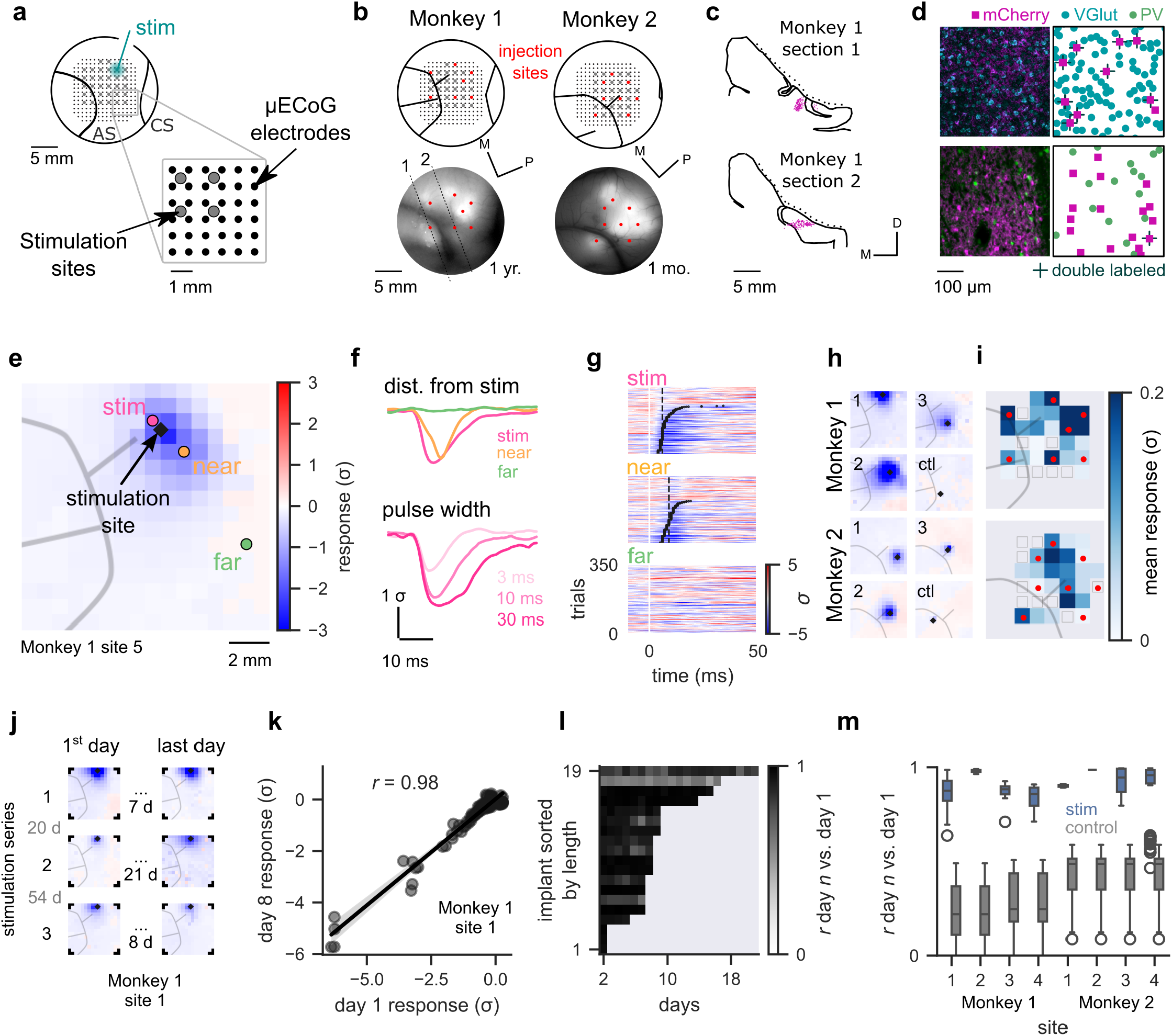
Combined optogenetics and µECoG revealed spatially specific, repeatable stimulation-evoked responses. **(a)** Illustration of µECoG electrode array and co-registered stimulation sites showing localization on the cortical surface (upper left) and details within the array (bottom right); AS arcuate sulcus, CS central sulcus. **(b)** Locations of injection sites in red overlaid on schematics of electrode and stimulation sites (top) and mCherry fluorescent signals (bottom). Dotted lines indicate coronal histological sections. **(c)** Locations of mCherry positive cells for the coronal sections in (b). Dotted lines indicate extent of recording chamber. **(d)** Histological images (left) and locations of identified cell bodies (right) for the two coronal sections in (b). Top row: stained with antibodies against mCherry (magenta) and in-situ hybridization against VGlut (cyan). Bottom row: stained with antibodies against mCherry (magenta) and parvalbumin (PV; green). **(e)** An example stimulation-evoked response in Monkey 1. Stimulation was delivered at the location marked with a diamond. **(f)** The evoked response over time as a function of distance from the stimulation site (top) and pulse width (bottom) for the example evoked response in Monkey 1. Colors correspond to the marked locations in (e). **(g)** Single trial evoked responses for the example in Monkey 1 at positions indicated in (e), sorted by response latency (black dots). Dashed lines indicate median latency. **(h)** Example stimulation-evoked responses across the cortical surface for 3 stimulation sites (numbered) and a control site far from viral expression (ctl). **(i)** Summary of the mean evoked response at each stimulation site. **(j)** Example of repeated stimulation to a single site over 3 array insertions (stimulation series), each spanning multiple days. **(k)** Pixelwise correlation between the response on a given day to the response on day 1, measured by Pearson correlation coefficient for the example stimulation site shown in (j). **(l)** Correlation coefficients between a given day and day 1 for repeated stimulation in 19 series across days. **(m)** Correlation coefficients from (l) organized by stimulation site and compared to control data with no stimulation (light grey). Box shows the median and quartiles; whiskers show range.

Optogenetic stimulation resulted in neural activity directly driven by ChR2 (*primary* responses), as well as neural activity putatively driven by connections with the neurons expressing ChR2 at the primary response site (*secondary* responses). Laser stimulation through one of the 32 holes in the µECoG array (a *stimulation site*) evoked changes in ECoG measurements across many electrodes (Fig 2e). Trial-averaged evoked potentials were typically larger close to the stimulation site and smaller at more distant electrodes (Monkey 1: linear regression response vs. distance *r*(7680) = 0.306, p < 0.001; Monkey 2: *r*(7680) = 0.321, p < 0.001) and varied in amplitude depending on the pulse width of stimulation (Monkey 1: linear regression response vs. pulse width *r*(11,204) = −0.137, p < 0.001; Monkey 2 *r*(19,130) = −0.417, p < 0.001; Fig 2f). Single-trial responses were more reliably driven near the stimulation site. We quantified single-trial reliability with the ratio of responsive trials to all trials (Monkey 1: linear regression response ratio vs. distance *r*(7680) = −0.108, p < 0.001; Monkey 2: *r*(7680) = −0.074, p < 0.001; Fig 2g). Across both animals, large responses across the µECoG array were observed following stimulation at some, but not all, stimulation sites (Fig 2h). We quantified the total size of a response by averaging the response across electrodes for each stimulation site. A summary of significant evoked responses for both animals (Fig 2i) revealed the strongest responses were located around the spur of the arcuate sulcus, closely resembling the pattern of AAV injections in both animals. Stimulation at sites far away from injections evoked little to no change in activity.

We leveraged our semi-chronic implants to take repeated measurements across days at select stimulation sites (a “stimulation series”). This revealed a consistent response evoked by stimulation during rest even after weeks had passed (Fig 2j). Each stimulation series lasted from two days to three weeks, with gaps of up to eight weeks in between measurements at the same site. We correlated the maximum evoked activity across electrodes from one day to the next to quantify stability (Fig 2k). For example, the correlation coefficient between day 8 and day 1 of site 1, series 1, in Monkey 1 was 0.98. We repeated this comparison for every day of a series to day 1 of that series, revealing stable evoked responses across many sites even for the longest series of 21 days (mean correlation coefficient across all series: 0.884, SD: 0.125; *N* = 21; Fig 2l). The average correlation coefficient at each site was significantly greater than when we repeated the analysis on data from the same days but with no optogenetic stimulation (Mann-Whitney U = 34694, n_1_ = 150, n_2_ = 232, p < 0.001; Fig 2m). We used a trial-adding analysis (see Methods) to test whether we collected sufficient stimulation data to expect stable network maps on each day. On average (N = 15 stimulation series with > 3,375 trials), correlation coefficients reached 90% of their maximum within 182 trials (SD 119 trials; Fig S1), well below the number of trials we recorded (350 for Monkey 1, 600 for Monkey 2).

Our combined stimulation and recording approach therefore had properties that enable large scale, longitudinal measurements of causal connectivity. Viral injections enabled optogenetic stimulation at distributed sites across motor cortices. Combining optical stimulation with electrical recordings let us measure across areas with minimal stimulation artifacts. Finally, evoked responses were spatially precise and consistent across days to weeks. We predicted that the spatial precision and repeatability of the evoked responses would therefore provide useful measurements of connectivity over multiple brain regions over time.

We used our stimulation and recording approach to assess analysis methods to estimate causal connectivity. We first explored analyzing the latency between stimulation and response, which is commonly used to distinguish primary and secondary responses to stimulation. The ratio of secondary response (the putative connections) to primary response (driven by stimulation) can then be used to estimate connectivity (SERR; Fig 3a). We calculated latency of stimulation-evoked responses in the high-gamma band to infer connectivity using SERR (see Methods: Latency-based connectivity estimation). Evoked responses had clear spatial patterns in their latency relative to stimulation, gradually increasing from 10 ms to 30 ms delays as distance from the stimulation location increases (Fig. 3b). We observed a distribution of latencies from electrodes with a detectable response to stimulation (Monkey 1: mean 21, SD 14 ms, *N* = 200; Monkey 2 mean 27, SD 18 ms, *N* = 106; Fig 3c). Across all sites, there was a positive relationship between median latency and distance from the stimulation site (Monkey 1: *R^2^* = 0.619, p < 0.001, n = 200; Monkey 2: *R^2^* = 0.618, p < 0.001, n = 106; Fig 3d).

**Figure 3.**
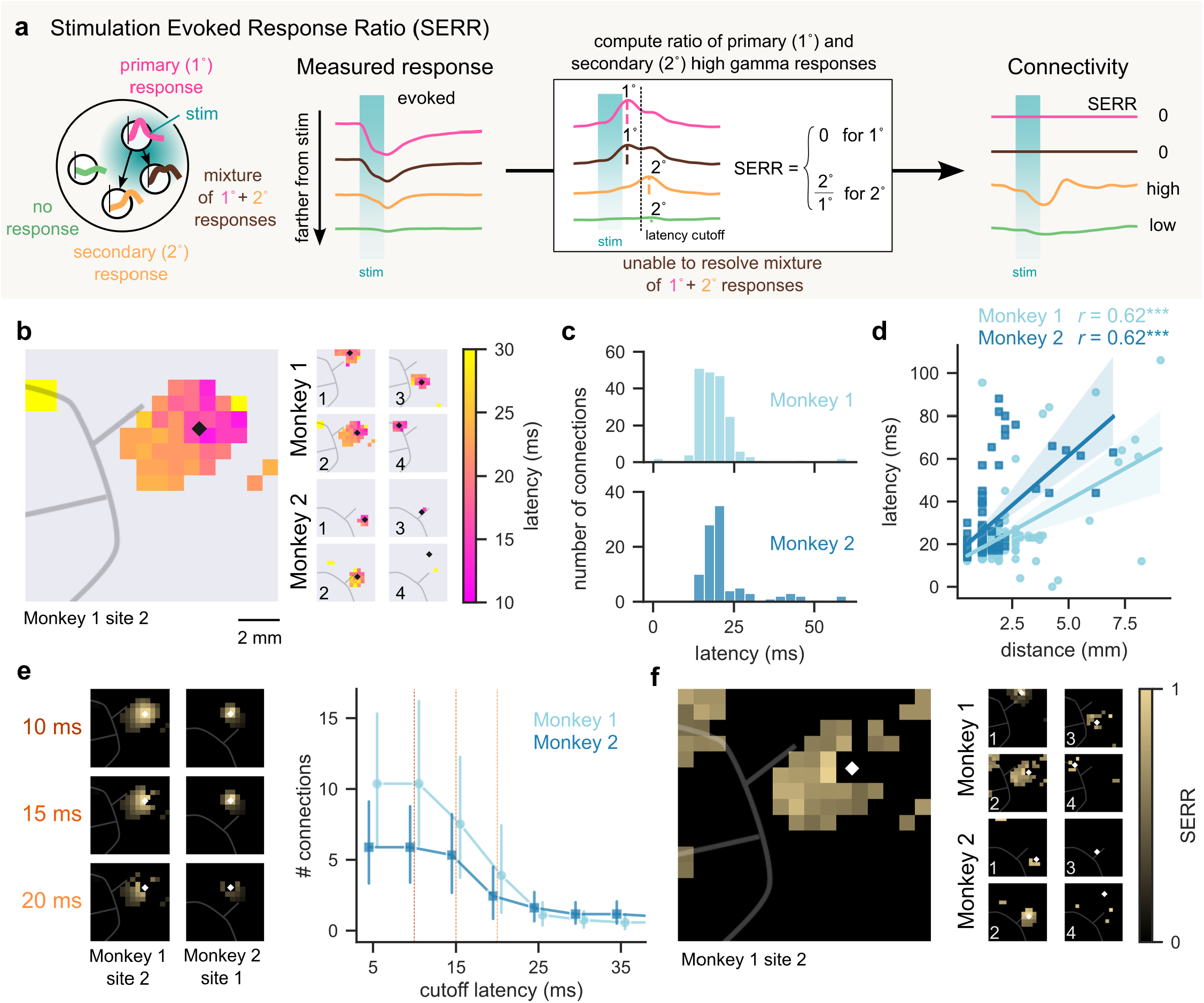
Stimulation-evoked response latency varies across space. **(a)** Illustration of connectivity estimation for an example network (left) using the stimulation evoked response ratio (SERR). High gamma (50 – 200 Hz) power was computed from stimulation-evoked responses (middle left), then divided into primary (1°, pink), secondary (2°, yellow), mixture of 1° and 2° (brown), and no (green) responses based on response amplitude and latency (middle right). Connectivity was computed as a ratio of 2° to 1° responses (right). **(b)** Spatial map of latency of detectable high gamma stimulation-evoked responses across the µECoG array for multiple example stimulation sites. **(c)** The number of significant connections identified as a function of latency across all sites for Monkey 1 (top, light blue) and Monkey 2 (bottom, dark blue). **(d)** Latency as a function of distance from the stimulation across all sites for Monkey 1 (light blue) and Monkey 2 (dark blue). Solid lines indicate linear regression; shading reflects a 95% confidence interval; *** indicates p < 0.001. **(e)** Example connectivity maps estimated with increasing latency cutoffs (left; top: 10 ms, middle: 15 ms, bottom: 20 ms); Mean number of estimated connections as a function of cutoff latency. Error bars reflect 95% confidence intervals. (N = 32 stimulation sites for Monkey 1, N = 18 for Monkey 2). **(f)** Example connectivity maps generated using SERR.

Classifying primary and secondary responses required choosing a cutoff latency, but this choice affected the resulting connectivity estimates. For example, as we increased the cutoff latency from 10 ms to 20 ms, the number of connections from sites 1 and 2 in monkey 1 decreased substantially (Fig 3e). In general, a lower selection for cutoff latency yielded more connections and a longer selection less. We saw modest evidence that the distribution of latencies was not unimodal (Monkey 1: Hartigans’ dip test statistic = 0.048, p = 0.002; Monkey 2: dip test statistic = 0.057, p = 0.013) but we did not find clear separation between “short” and “long” latency responses. Example maps of SERR connectivity (18 ms cutoff) show that latency-based maps were often sparse, related to binarizing the data into primary and secondary responses (Fig. 2f). However, capturing significantly delayed responses to stimulation demonstrated our ability to measure causal connectivity across brain regions. We hypothesized that other analysis methods that did not rely on limited binary classification may be able to resolve higher resolution maps of connectivity.

To better resolve primary and secondary responses, we developed a stimulation-locked imaginary coherence (SLIC) metric that detects directed connectivity via the phase of responses (Fig 4a). We calculated SLIC using the imaginary coherence (Nolte et al., 2004) between each electrode and the stimulation site time-aligned to stimulation onset. Using imaginary coherence allowed us to remove components of the measured coherence with zero time-lag (see illustration). When the phase difference between the stimulation site and a given electrode is zero, then imaginary coherence is zero. Imaginary coherence is nonzero when the two signals are coherent and out of phase. Applying SLIC to data aligned to stimulation therefore should only detect temporally delayed coherence, thereby improving identification of secondary, rather than direct, responses to stimulation input. We found that the average phase difference between the stimulation site and each electrode revealed consistent nonzero phase difference around the stimulation, consistent with the existence of secondary responses that could be measured by their phase (Fig. 4b).

**Figure 4.**
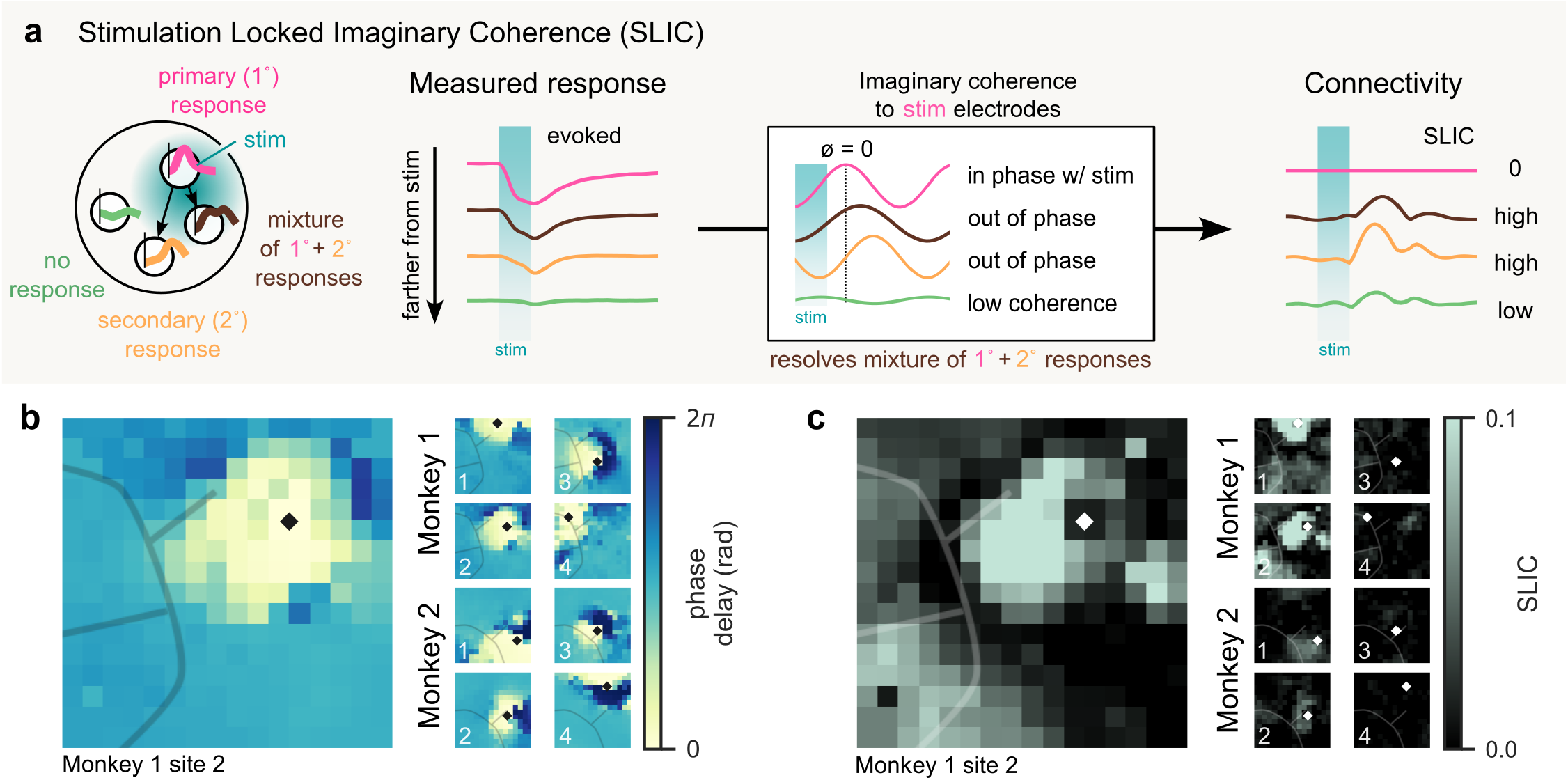
Phase-based analysis reveals secondary responses. **(a)** Illustration of connectivity estimation for an example network (left) using stimulation locked imaginary coherence (SLIC). Stimulation-evoked responses were computed (middle left), and imaginary coherence on the evoked response was computed between each electrode and the electrodes nearest to the stimulation site (middle right), which was used to calculate SLIC-based connectivity (right). When these signals are coherent and in phase, SLIC connectivity is low (electrodes with primary responses only; pink). If they are coherent and out of phase, SLIC connectivity is high (electrodes with any secondary response; yellow and brown). Finally, if the coherence between signals is low, SLIC connectivity is also low (electrodes with no response; green). **(b)** Spatial maps of phase delays from the stimulation site to each electrode for multiple example stimulation sites. Note that two random signals will have a phase delay of π in this analysis. **(c)** Spatial maps of SLIC connectivity estimates for the example stimulation sites shown in (b).

There was a striking similarity between the phase maps (Fig 4b) and the latency maps (Fig 3b), yet phase differences could be estimated everywhere while latency was only calculated at electrodes with detected responses in high gamma. SLIC, correspondingly, produced maps that could detect and quantify statistically significant connectivity across the full span of the array (Fig 4c). Patterns of connectivity measured with SLIC were qualitatively similar to those measured using latency-based SERR (see Fig 3f), but with more complete spatial sampling since there was no explicit removal of electrodes with primary responses. This highlights that SLIC could address challenges of mesoscale connectivity analyses, such as resolving secondary responses that may be mixed with primary responses close to the site of stimulation. We next sought to quantify differences between latency and phase-based analyses and compare SLIC connectivity to established activity-based methods of functional connectivity.

Our SLIC metric yielded maps of connectivity qualitatively similar to those we calculated using SERR. We quantified how many connections were captured by these two causal metrics and a common functional connectivity metric to compare their sensitivity. Using SLIC, we observed many statistically significant connections (see Methods) across the 32 stimulation sites (Monkey 1: mean 74, SD 41 connections per site; Monkey 2: mean 51, SD 35 per site; across 240 electrodes). SERR captured a comparatively modest network of connections (Monkey 1: mean 5, SD 9 connections per site; Monkey 2: mean 3, SD 5). We used granger prediction (GP) to estimate the functional connectivity between each channel applied to resting state data with no stimulation and to stimulation data (stimulation evoked Granger prediction, SEGP; see Methods). Both GP and SEGP captured many significant connections (GP - Monkey 1: mean 192, SD 22 connections per site; Monkey 2: mean 158, SD 30; SEGP - Monkey 1: mean 190, SD 33; Monkey 2: mean 128, SD 45). Applying SEGP recovered a similar number of connections as GP (Monkey 1: Wilcoxon signed-rank test Z = 152, p = 0.04; Monkey 2: Z = 228, p = 0.51), suggesting that the number of identified connections was not necessarily solely influenced by providing a causal network input (i.e. passive functional connectivity versus causal connectivity). On the other hand, applying latency-based SERR instead of SLIC analyses to the same stimulation data revealed significantly fewer connections (Monkey 1: Wilcoxon signed-rank test Z = 0.0, p < 0.001; Monkey 2: Z = 0.0, p < 0.001). We therefore conclude that the analysis method choice substantially affected the ability to detect connections.

We next investigated the spatial similarity of connectivity computed across different metrics. Measurements that were driven by stimulation were visually similar to one another (left panels of Fig 5a). We chose to compare SLIC, SEGP, and GP since they detected similar numbers of connections compared to the more conservative SERR metric. We repeatedly computed SLIC, SEGP, and GP on two random splits of data (Fig 5b). For each random split, we computed the correlation between all six computed maps of connectivity (two maps for each metric; Fig 5c). We expected that maps produced by the same method would be consistent, whereas comparisons across methods would be less correlated. In addition, we expected both maps generated using stimulation to be more similar than maps generated with different underlying data. Between SLIC and GP, within-method correlation coefficients were always higher than across-method coefficients, suggesting that the two methods produced different connectivity maps (Fig 5d). Comparisons of Granger analysis applied to rest or stimulation-evoked activity (GP and SEGP, respectively) showed that the within- and across-method correlation coefficients were often similar to those of within-method, suggesting that the two methods produced similar connectivity maps (Fig 5e). While we found that analysis method played a larger role than the presence of stimulation for detecting connections (Fig. 4), the magnitude of the response evoked by stimulation may influence the ability to reliably estimate connections. Consistent with this, we found that SLIC and Granger generally identified more similar connectivity patterns when applied to stimulation data as opposed to resting data (SLIC-SEGP > SLIC-GP, Wilcoxon signed-rank test, Z = 403, p < 0.001, N = 28). Moreover, SLIC and SEGP found more similar connectivity patterns when stimulation evoked larger responses in the network (Fig. 5f).

**Figure 5.**
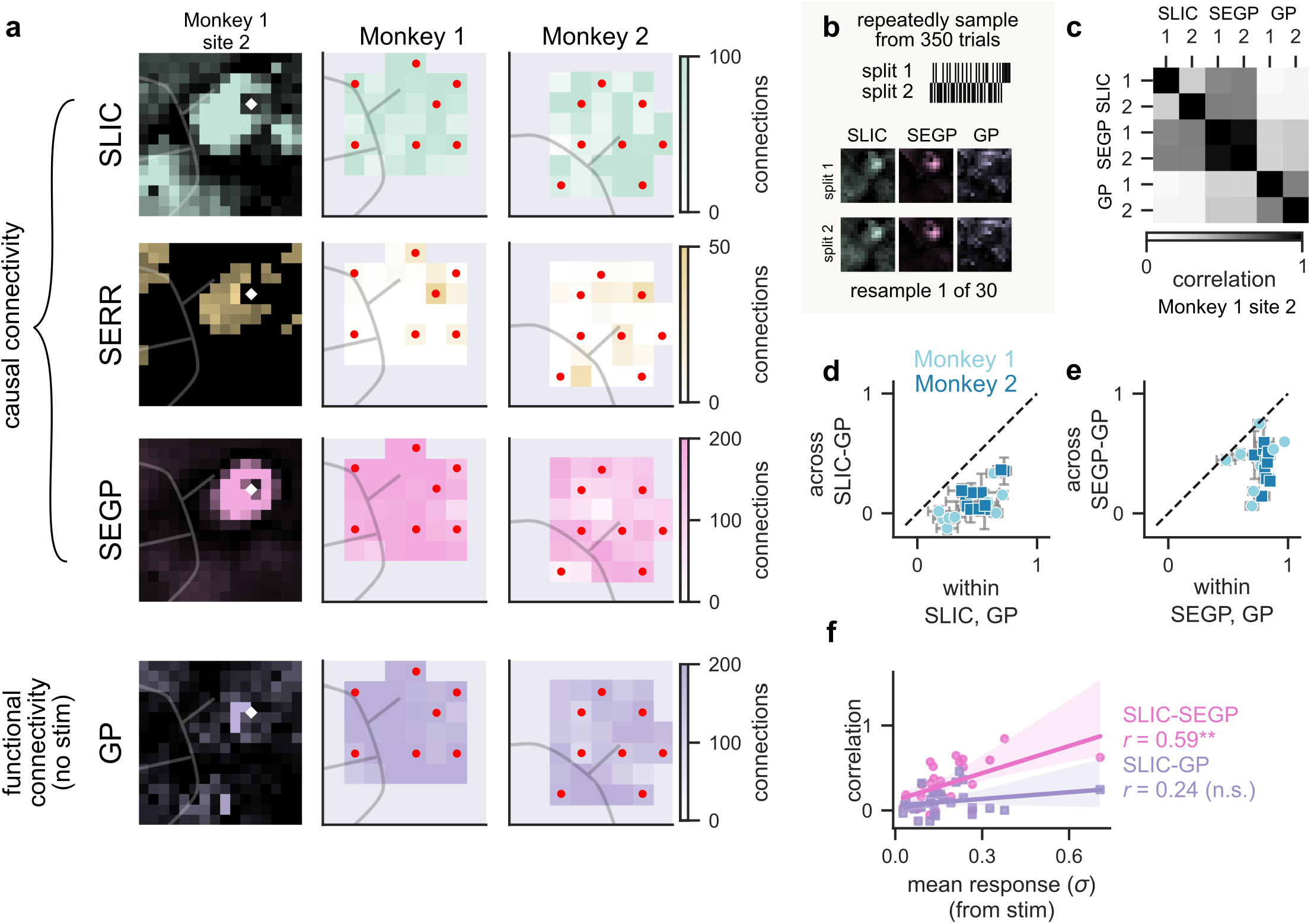
Comparison of connectivity metrics. **(a)** Comparison of causal and functional connectivity metrics applied to single day µECoG data (rows, from top to bottom: SLIC, SERR, SEGP, and GP). Left column: spatial maps of connectivity generated using different metrics for an example stimulation site. Right columns: summary of the number of significant connections identified from each stimulation site (source for functional connectivity) for Monkey 1 (middle) and Monkey 2 (right). **(b)** Illustration of calculations to directly compare the spatial pattern of connectivity across different metrics. We repeatedly divided trials into two randomized splits (top) and then computed connectivity using all metrics for each split (bottom), and computed correlation between all map pairs. **(c)** The correlation coefficients across pixelwise comparisons within and across each metric for one example stimulation site (Monkey 1, site 2; shown in (a)). **(d)** Correlations between across metric (SLIC to GP) maps as a function of correlations within-metric (SLIC, GP) for all stimulation sites with a significant response (Monkey 1, light blue; N = 17; Monkey 2, dark blue, N = 11). Error bars reflect standard deviation from the mean. Dashed black line reflects the unity line. **(e)** Same as (d), but comparing GP to SEGP. **(f)** Relationship between across-metric correlation and mean response from stimulation for SLIC-GP (purple) and SLIC-SEGP (pink) comparisons across all stimulation sites with a significant response (N = 28). Solid lines show linear regression; shading reflects a 95% confidence interval. ** indicates p < 0.01. n.s. not significant.

The ability to capture physiologically meaningful connections is an important feature of any metric. Although we don’t know the ground truth connectivity, we estimate that there are anatomical connections underlying a vast majority of electrode pairs in our setup based on the ubiquity of cortico-cortical connections in motor cortices (Gatter et al., 1978; Huntley & Jones, 1991; Ninomiya et al., 2019). Latency-based SERR detected few connections, suggesting that this metric may not reliably detect connections and likely underestimates network connectivity. Our results also show that causal connectivity maps, measured using stimulation, provide different estimates of network connectivity, and that large causal stimulation input can reveal similar network structures regardless of analysis method (Granger vs. SLIC).

We examined the frequency-specific components of connectivity to better compare the spatiotemporal properties of each metric. Cortical circuitry can give rise to activity across multiple frequency bands, each thought to reflect circuit-specific computations (MacKay, 2004), suggesting that spectral analyses like SLIC and Granger may also be able to resolve frequency-specific networks. We computed maps of connectivity for SLIC, GP, and SEGP across five frequency bands: 0.5-12, 12-30, 30-80, 80-120, and 120-200 Hz (Fig 6a). Qualitatively, SLIC maps that appeared more distinct across frequencies compared to the other approaches, often revealing long-range connections even at high frequencies. In contrast, GP maps revealed more local connectivity and were visually similar across frequencies. We quantified these differences across bands by performing pairwise correlations across maps computed in different frequency bands (Fig 6b). We found more distinct connectivity maps across frequencies using SLIC (Monkey 1: mean correlation 0.36, SD 0.35, *N* = 17 stimulation sites with significant responses; Monkey 2: mean 0.38, SD 0.36, *N =* 11) than using either GP (Monkey 1: mean correlation 0.61, SD 0.24, *N* = 17; Monkey 2: mean 0.68, SD 0.21, *N =* 11) or SEGP (Monkey 1: mean correlation 0.77, SD 0.15, *N* = 17; Monkey 2: mean 0.77, SD 0.19, *N =* 11). Because SEGP and SLIC were computed on the same stimulation data, these results make it apparent that the inherently time-dependent Granger method lacks the frequency resolution of SLIC.

**Figure 6.**
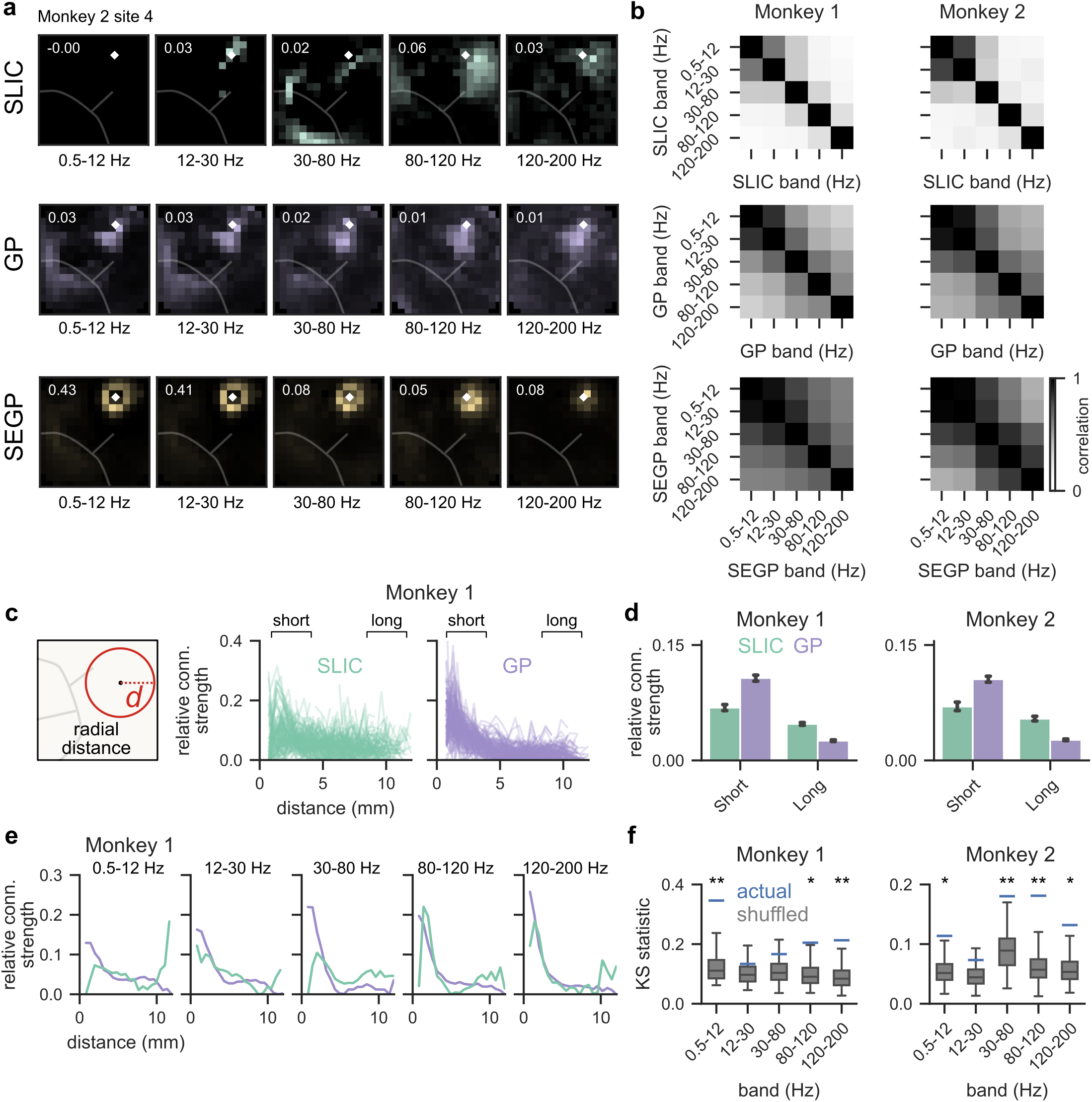
Frequency and spatial specificity of causal and functional connectivity measurements. **(a)** Spatial maps of connectivity for one example stimulation site (Monkey 2, site 4) calculated with different analysis metrics (rows), separated by frequency band (columns). For each map, colors are scaled from zero to an upper limit indicated by the inset text. **(b)** Matrix of the average correlation between connectivity maps at different frequency bands for different analysis metrics (rows) and monkey (columns; Monkey 1: N = 17 stimulation sites; Monkey 2: N = 11). **(c)** Left: illustration of radial distance between electrode pairs (red circle) overlayed on the cortical surface. Right: Relative connection strength as a function of radial distance computed with SLIC (middle right) and GP (far right). Each line reflects a different stimulation site (Monkey 1 only). **(d)** Relative strength of short-range (less than 4 mm) and long-range (greater than 8 mm) connections for SLIC (green) and GP (purple). Bars indicate mean; error bars denote 95% confidence intervals (N = 32 stimulation sites). Distance cutoffs are indicated in (c). **(e)** Relative connection strength as a function of radial distance pooled across all stimulation sites for Monkey 1, separated by frequency band (columns). **(f)** KS difference between SLIC and GP distributions for all stimulation sites across frequency bands. Box shows the median and quartiles; whiskers show range (N = 32 stimulation sites). * indicates p < 0.05, ** p < 0.01.

We analyzed the distance-dependent decay of connection strength to quantify the ability of each metric to resolve both local and long-range connections. We focused this analysis on SLIC and GP to highlight the advantages of our causal method over functional connectivity. We first calculated the relative connection strength at a given radius from each stimulation site individually (Fig 6c). Qualitatively, the distributions of connectivity appeared more varied across distance for SLIC connectivity compared to GP, which was mostly concentrated at short distances. We quantified the relative strength of long-range connections at distances above 8 mm from the stimulation site (Fig 6d). Across frequency bands, long-range SLIC connectivity was significantly greater than long-range GP connectivity (Monkey 1: Mann-Whitney U = 220199, n_1_ = 549, n_2_ = 646, p < 0.001; Monkey 2: U = 194956, n_1_ = 460, n_2_ = 650, p < 0.001). In contrast, local SLIC connectivity (less than 4 mm from stimulation) was significantly less than local GP connectivity (Monkey 1: Mann-Whitney U = 783327, n_1_ = 1,167, n_2_ = 1,188, p < 0.001; Monkey 2: U = 696497, n_1_ = 1,167, n_2_ = 1,225, p < 0.001).

The above results suggested that each method captured distinct spatial scales of connections. We compared the difference in spatial distributions of SLIC and GP connectivity (across all stimulation sites) using the Kolmogorov-Smirnov (KS) distance (Fig 6e). The majority of the five frequency bands we analyzed (Monkey 1: three of five; Monkey 2: four of five) had significantly higher KS distance between SLIC and GP distributions compared to shuffled distributions (p < 0.05 based on 1000 random permutations; Fig 6f). This confirms that SLIC and GP differ in their ability to detect local-versus long-range connections. These differences may be one significant source of variability in networks revealed by functional versus causal connectivity methods (Fig.5).

Single measurements of connectivity like the ones presented so far only capture a snapshot of the measured network. Longitudinal measurements will improve our ability to characterize network architectures, their changes over time, and relationships between networks and behavior. We compared connectivity across longitudinal stimulation series to test the stability of our measurements across long periods of time (Fig 7a). SLIC connectivity was significantly more consistent than null data with no stimulation in seven out of eight sites we tested (Mann-Whitney U test, p < 0.05; Table S1; Fig 7b). In contrast, GP was no more consistent across days than time-shuffled data in six of the eight sites we tested (Mann-Whitney U test, p > 0.05; Table S1; Fig 7c). The consistency across days at each site was positively correlated with the size of the evoked response to stimulation for SLIC causal connectivity, but not for GP functional connectivity (Fig 7d). In other words, connectivity measurements were most consistent across days at stimulation sites where we had the ability to drive the network strongly. Under the assumption that networks remain similar across time in the absence of significant drivers of change (e.g., learning), this further demonstrates advantages of causal methods over functional connectivity.

**Figure 7.**
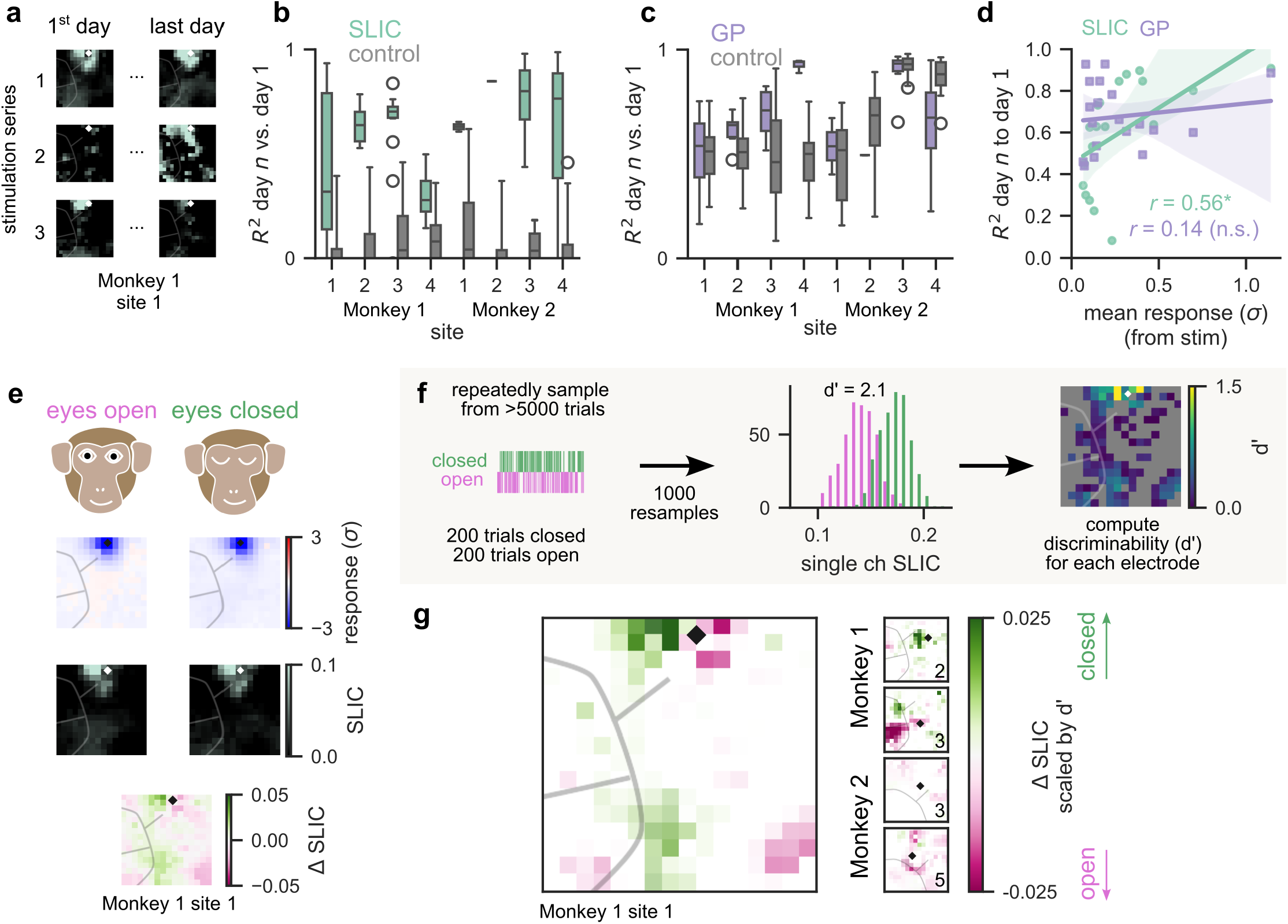
Stimulation-based connectivity stable across days to weeks but sensitive to changes in behavioral state. **(a)** Example of repeated SLIC connectivity maps for a single stimulation site over 3 array insertions (stimulation series), each spanning multiple days. **(b)** Correlation between connectivity maps on each day with that of day 1 for repeated SLIC measurements (green) compared to control (grey; SLIC with no stimulation) for multiple stimulation sites in each monkey. Box indicates the mean and quartiles; whiskers indicate range. **(c)** Same as (b) but with GP connectivity (control reflects time-shuffled data). **(d)** Relationship between across-day connectivity map correlation and the mean response to stimulation for each stimulation site. Solid lines reflect linear regression; shading indicates 95% confidence interval. **(e)** Example spatial maps of broadband responses (upper middle) and SLIC connectivity (lower middle) on stimulation trials where animals had their eyes open (left) or closed (right), and the map of the change in SLIC between the two behavior states (bottom). **(f)** Illustration of the resampling procedure to estimate differences in SLIC. For each electrode, we generated two distributions of 1000 SLIC values for eyes open and eyes closed trials drawn randomly with replacement (left). We then calculated discriminability (d’) between those distributions (middle) and tested for significance against 50 values of d’ computed with shuffled labels. **(g)** Spatial map of the difference between SLIC connectivity computed in the eyes closed versus eye open state, scaled by d’ for multiple example stimulation sites across both monkeys.

The prior analysis shows that we can consistently estimate connectivity over days under settings where we expect minimal changes, but a meaningful connectivity metric should also be able to detect functionally relevant changes in networks. For instance, latency-based causal connectivity has been shown to vary trial to trial in state-dependent ways (Qiao et al., 2020). If SLIC measurements solely reflect fixed anatomical pathways, then we would expect our measurements to be stable across behavioral states. But if SLIC also captures functional changes in connectivity strength, then different behaviors might lead to differences in causal connectivity. We kept track of the position of the animals’ eyes during recording, including whether they were open or closed. We hypothesized that open and closed eye behavior states may correspond to different patterns of connectivity. When we divided our data into trials in which animals’ eyes were open or closed, we observed qualitatively similar stimulation-evoked broadband responses, but noticeable differences in connectivity measured using SLIC (Fig 7e). To test whether there were any detectable differences in connectivity between these two behavioral states, we adopted a re-sampling approach to estimate confidence bounds for each connectivity estimate (Fig 7f). For each electrode, we generated a distribution of connectivity measurements drawn from random subsets of eyes open and eyes closed trials. For each distribution we measured the separability of the two behavioral states using d’, which measures distance as a factor of the pooled standard deviation of the distributions (Green & Swets, 1988; Rouse et al., 2013). For comparison, we also computed d-prime between distributions generated from shuffling the eye state labels. Across five stimulation sites, we observed significant differences in SLIC connectivity in some, but not all, electrodes (Mean across sites 125/240 electrodes, SD 27 electrodes with p < 0.01; Wilcoxon signed rank test, false discovery rate corrected; Fig 7g). These electrodes tended to be clustered around the stimulation site, with some smaller differences detectable at longer distances. Thus, SLIC measurements were sensitive to the changes in behavior between open and closed eye states. Taken together, the stability of our measurements at rest and the difference between connectivity measured across states suggests that our approach is well-suited to map behaviorally relevant connectivity changes in mesoscale networks longitudinally.

## Discussion

Evaluating causal network connections across large portions of the brain over time requires new methods to repeatedly deliver stimulation and new analyses to separate stimulation-driven and secondary responses. We addressed these challenges with simultaneous optogenetic stimulation and µECoG recordings in awake macaques combined with phase-based connectivity analysis. Our causal measurements were stable across days to months, demonstrating feasibility to monitor cortical connectivity over timeframes relevant for behavior. The stimulation-locked imaginary coherence (SLIC) metric that we developed captured frequency-specific and spatially heterogenous network properties that functional connectivity methods failed to consistently detect, and had the sensitivity to detect significant state-dependent differences. Our combined methodological and analytical approach for making causal measurements of connectivity therefore enables spatially localized, longitudinal characterization of large brain networks during behavior.

### Advantages of causal connectivity

Our comparisons between causal and functional connectivity highlighted that causal (stimulation-based) maps were similar regardless of the analysis metric, but distinct from functional (resting-based) maps. We observed significantly different connectivity maps when applying the same Granger analysis to resting activity (GP) versus stimulation evoked activity (SEGP; Fig 5). Further, SEGP and SLIC maps were more similar when stimulation evoked large neural responses, suggesting that stimulation drove a distinct network from what could be detected at rest, regardless of the analytical approach used to identify connections. Relying on ongoing activity to estimate connections appears unable to capture this driven network, which may reflect a meaningful route of flexible neural communication (Daie et al., 2026; Qiao et al., 2020). Our results are consistent with prior observations that different networks were estimated using stimulation compared to resting activity alone (Bauer et al., 2018; Finkelstein et al., 2025). Our study reinforces the advantages of including causal connectivity measures to more fully capture connectivity and extend its application to large networks in primates.

### Analytical challenges using mesoscale stimulation and recording

Mesoscale recordings like ECoG raise concerns of potential spatial resolution limitations, for instance due to passive volume conduction. Our approach leveraged arrays that provided spatial resolutions on the millimeter scale (Trumpis et al., 2021), consistent with a growing body of studies showing that high density µECoG can provide high spatiotemporal resolution maps of cortical activity (Kaiju et al., 2017; Li et al., 2023; Ouchi et al., 2025; Wang et al., 2017). Indeed, we found connectivity patterns that had non-gaussian spatial distributions (Fig 5) and did not strictly decay with distance (Fig 6), neither of which would be predicted by passive volume conduction. Our approach identified many long-range connections, consistent with anatomical tracer experiments (Gatter et al., 1978; Huntley & Jones, 1991; Ninomiya et al., 2019). This suggests that our causal measurements resolved circuit architecture rather than sensor-level artifacts.

Our systematic comparison of analysis methods demonstrated that algorithm choice strongly impacts estimates of causal connectivity from µECoG signals. We initially used a latency-based approach (Fig 3) that has been successful when neural activity was measured far from the stimulation site (Qiao et al., 2020; Yazdan-Shahmorad et al., 2018). We found that using a binary threshold to separate connected-from stimulation-driven activity was too coarse, reducing detection sensitivity (Fig 5). This issue likely arises from the mixture of primary and secondary responses in mesoscale signals like µECoG (Fig 1b). Our results suggest that coherence analyses are better suited to mesoscale signals because they can leverage phase information to more directly resolve secondary responses even when mixed with primary, stimulation-driven activity (Fig 4).

Different frequency bands are thought to reflect activity patterns of distinct microcircuits (MacKay, 2004), suggesting that band-specific connectivity estimates could be helpful to distinguish microcircuit-specific network interactions. Our SLIC measurements resolved band-specific differences in connectivity patterns that were not detected with SEGP analyses on the same data (Fig 6a,b). The inherent frequency dependence of SLIC’s coherence analysis, in contrast with time domain autoregression used in Granger analyses, likely underly this improved frequency resolution. We acknowledge that we violated nonstationarity assumptions by applying Granger methods to stimulation-evoked data but included this analysis to demonstrate the implications that algorithmic differences have on connectivity analysis. Only our approach using SLIC captured frequency-specific components of connectivity, allowing discrimination between microcircuits. Pairing SLIC frequency-specific connectivity dissection with stimulation patterns designed to drive specific microcircuits (Sohal et al., 2009) will provide paths to further refine connectivity estimation methods and dissect network changes during behavior.

### Longitudinal stability and state dependence

Our work addresses gaps in tools to monitor network connectivity over time during behavior. While longitudinal measurements are subject to various sources of noise – for example, shifts in the position of the electrodes, changes in the fiberoptic coupling, and changes in surface opacity – our measurements of causal connectivity remained remarkably stable over repeated measurements spanning weeks to months (Fig 7). This stability demonstrates that our approach can enable new experiments to quantify connectivity changes across both short (days) and long (weeks, months) timeframes relevant to behaviors like motor learning. Stability across daily repeated measurements might suggest that our assay reflects mesoscale anatomical connectivity that does not typically change substantially over time. However, the significant differences we saw between behavioral states indicate that our approach is sensitive to rapid reconfigurations of neural circuits thought to occur during learning (Ogawa et al., 2022; Perich et al., 2018). The ability to detect these rapid changes suggest that our causal measurements may capture a mixture of more slowly varying anatomical connectivity and dynamic, functional connections within neural networks.

### Limitations and future work

Having resolved many challenges inherent to mesoscale signals, our approach establishes a foundation for large-scale causal connectivity mapping. We focused our initial experiments on spatially localized injections to control the overlap of stimulated regions. Our present study confirmed that SLIC can resolve these overlaps, which opens the opportunity to extend our technique to more densely map network connectivity using more global channelrhodopsin expression (Yazdan-Shahmorad et al., 2016). Algorithmic advances would also improve measurement efficiency, which will be critical to increasing the size of networks that can be mapped (Grosenick et al., 2015). For example, we used a fixed number of stimulation trials for all measurements, but dividing our data into groups of increasing size revealed that some stimulation sites required fewer trials to achieve a consistent map (Fig S1). Future work could further optimize the time needed to estimate connectivity from a given stimulation site using closed-loop algorithms (Draelos et al., 2025; Lepage et al., 2013; Ouchi & Orsborn, 2022). Moreover, while we focus on measuring connectivity, our longitudinal mesoscale recording and stimulation technique also provides new experimental opportunities to probe distributed neural computations. For example, connectivity measurements could be paired with optogenetic stimulation to experimentally test the robustness of learned behaviors or down-stream neural representations to perturbations. Bridging the gap between static anatomy and dynamic behavior requires tools that can reliably track causal interactions. Our approach provides the necessary foundation to map these dynamic, distributed computations in the behaving primate.

## Acknowledgements

We would like to thank Jonathan Viventi and colleagues for providing the electrode arrays used in this work, and Greg Horwitz and colleagues for assistance with histological procedures. This work was supported in part by a postdoctoral fellow award from Weill Neurohub (L.R.S.), a National Center for Advancing Translational Sciences of the National Institutes of Health fellowship (TL1 TR002318, R.A.C.), an NSF Accelnet INBIC fellowship (P.R.), a Simons Collaboration for the Global Brain Pilot award (898220, A.L.O.), and the National Institute of Neurological Disorders and Stroke (NIH R01 NS134634, A.L.O.).

## Supplemental Figures and Tables

**Figure S1.**
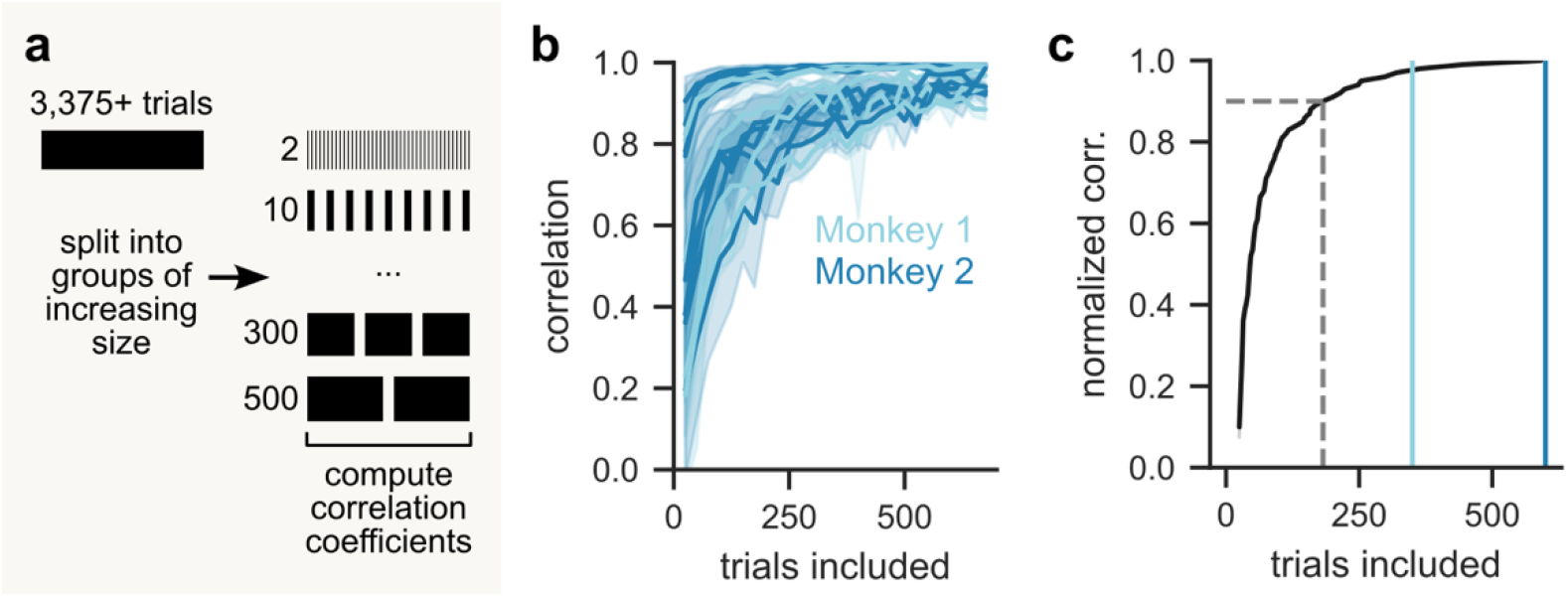
Trial adding analysis. **(a)** Illustration of analyses. We split stimulation datasets containing many trials (right) into groups of increasing size (rows depicted on the left) and then calculated the correlation between evoked at each size. **(b)** The correlation between pairs of evoked response maps as a function of the number of trials used to estimate the map. Each line is a different stimulation series. Solid lines indicate the mean; shaded regions indicate standard deviation away from the mean across multiple pairs of the same trail size (*N* = 15 series). **(c)** Correlation from (b) was normalized so that the maximum was always 1.0. The mean normalized correlation across series is plotted against number of trials (*N* = 15). Gray dashed line indicates trials needed for 90% of the maximum correlation. Solid lines indicate the number of trials recorded each day for Monkey 1 (light blue) and Monkey 2 (dark blue).

---

**Table S1.**
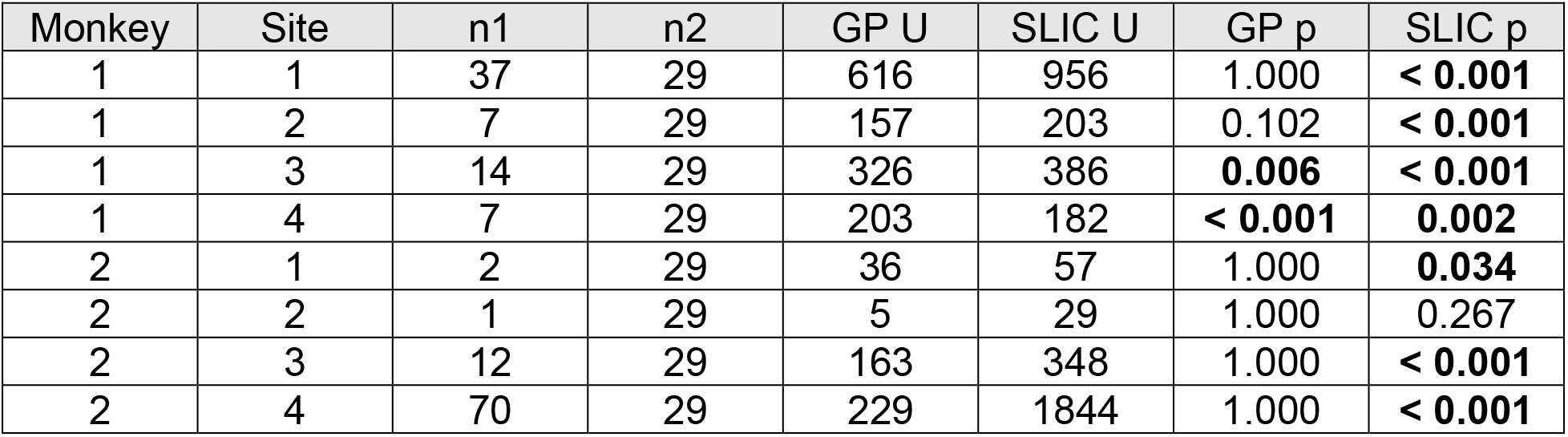
Longitudinal statistics information for all stimulation sites across both monkeys. For SLIC, n1 and n2 represent the sample sizes (number of days) for stimulation data and resting data, respectively. For GP, n1 and n2 represent the sample sizes for actual data and shuffled data, respectively. P-values less than 0.001 are reported as < 0.001. P-values were Bonferroni corrected for multiple comparisons; adjusted values mathematically exceeding 1 are reported as 1.000.

| Monkey | Site | n1 | n2 | GP U | SLIC U | GP p | SLIC p |
| --- | --- | --- | --- | --- | --- | --- | --- |
| 1 | 1 | 37 | 29 | 616 | 956 | 1.000 | < 0.001 |
| 1 | 2 | 7 | 29 | 157 | 203 | 0.102 | < 0.001 |
| 1 | 3 | 14 | 29 | 326 | 386 | 0.006 | < 0.001 |
| 1 | 4 | 7 | 29 | 203 | 182 | < 0.001 | 0.002 |
| 2 | 1 | 2 | 29 | 36 | 57 | 1.000 | 0.034 |
| 2 | 2 | 1 | 29 | 5 | 29 | 1.000 | 0.267 |
| 2 | 3 | 12 | 29 | 163 | 348 | 1.000 | < 0.001 |
| 2 | 4 | 70 | 29 | 229 | 1844 | 1.000 | < 0.001 |

## Notes

### Competing Interest Statement

A.L.O. is a scientific advisor for Meta Reality Labs and a consultant for EpiaNeuro. All other authors declare no competing interests.

## References

Avery, M. C., & Krichmar, J. L. (2017). Neuromodulatory Systems and Their Interactions: A Review of Models, Theories, and Experiments. Frontiers in Neural Circuits, 11. 10.3389/fncir.2017.00108

Balbinot, G., Milosevic, M., Morshead, C. M., Iwasa, S. N., Zariffa, J., Milosevic, L., Valiante, T. A., Hoffer, J. A., & Popovic, M. R. (2025). The mechanisms of electrical neuromodulation. The Journal of Physiology, 603(2), 247–284. 10.1113/JP286205

Banerjee, A., Dean, H. L., & Pesaran, B. (2010). A Likelihood Method for Computing Selection Times in Spiking and Local Field Potential Activity. Journal of Neurophysiology, 104(6), 3705–3720. 10.1152/jn.00036.2010

Bauer, A. Q., Kraft, A. W., Baxter, G. A., Wright, P. W., Reisman, M. D., Bice, A. R., Park, J. J., Bruchas, M. R., Snyder, A. Z., Lee, J.-M., & Culver, J. P. (2018). Effective Connectivity Measured Using Optogenetically Evoked Hemodynamic Signals Exhibits Topography Distinct from Resting State Functional Connectivity in the Mouse. Cerebral Cortex (New York, NY), 28(1), 370–386. 10.1093/cercor/bhx298

Bundy, D. T., Barbay, S., Hudson, H. M., Frost, S. B., Nudo, R. J., & Guggenmos, D. J. (2023). Stimulation-Evoked Effective Connectivity (SEEC): An in-vivo approach for defining mesoscale corticocortical connectivity. Journal of Neuroscience Methods, 384, 109767. 10.1016/j.jneumeth.2022.109767

Buzsáki, G., Anastassiou, C. A., & Koch, C. (2012). The origin of extracellular fields and currents—EEG, ECoG, LFP and spikes. Nature Reviews. Neuroscience, 13(6), 407–420. 10.1038/nrn3241

Chiang, C.-H., Won, S. M., Orsborn, A. L., Yu, K. J., Trumpis, M., Bent, B., Wang, C., Xue, Y., Min, S., Woods, V., Yu, C., Kim, B. H., Kim, S. B., Huq, R., Li, J., Seo, K. J., Vitale, F., Richardson, A., Fang, H., … Viventi, J. (2020). Development of a neural interface for high-definition, long-term recording in rodents and nonhuman primates. Science Translational Medicine, 12(538), eaay4682. 10.1126/scitranslmed.aay4682

Cho, Y. U., Lim, S. L., Hong, J.-H., & Yu, K. J. (2022). Transparent neural implantable devices: A comprehensive review of challenges and progress. Npj Flexible Electronics, 6(1), 53. 10.1038/s41528-022-00178-4

Cohen, M. X. (2014). Analyzing Neural Time Series Data: Theory and Practice. The MIT Press. 10.7551/mitpress/9609.001.0001

Daie, K., Aitken, K., Rózsa, M., Bull, M. S., Humphreys, P. C., Wang, Z. C., Kinsey, L., Kulkarni, M., Stachenfeld, K. L., Eckstein, M. K., Kurth-Nelson, Z., Clopath, C., Lillicrap, T. P., Botvinick, M., Golub, M., Mihalas, S., & Svoboda, K. (2026). Functional reorganization of motor cortex connectivity during learning. bioRxiv, 2026.03.03.709199. 10.64898/2026.03.03.709199

Das, A., & Fiete, I. R. (2020). Systematic errors in connectivity inferred from activity in strongly recurrent networks. Nature Neuroscience, 23(10), 1286–1296. 10.1038/s41593-020-0699-2

Denovellis, E. L., Myroshnychenko, M., Sarmashghi, M., & Stephen, E. P. (2022). Spectral Connectivity: A python package for computing multitaper spectral estimates and frequency-domain brain connectivity measures on the CPU and GPU. Journal of Open Source Software, 7(80), 4840. 10.21105/joss.04840

Draelos, A., Loring, M. D., Nikitchenko, M., Sriworarat, C., Gupta, P., Sprague, D. Y., Pnevmatikakis, E., Giovannucci, A., Benster, T., Deisseroth, K., Pearson, J. M., & Naumann, E. A. (2025). A software platform for real-time and adaptive neuroscience experiments. Nature Communications, 16(1), 9909. 10.1038/s41467-025-64856-3

Feng, G., Jensen, F. E., Greely, H. T., Okano, H., Treue, S., Roberts, A. C., Fox, J. G., Caddick, S., Poo, M., Newsome, W. T., & Morrison, J. H. (2020). Opportunities and limitations of genetically modified nonhuman primate models for neuroscience research. Proceedings of the National Academy of Sciences, 117(39), 24022–24031. 10.1073/pnas.2006515117

Finkelstein, A., Daie, K., Rózsa, M., Darshan, R., & Svoboda, K. (2025). Connectivity underlying motor cortex activity during goal-directed behaviour. Nature, 1–7. 10.1038/s41586-025-09758-6

Gatter, K. C., Sloper, J. J., & Powell, T. P. (1978). The intrinsic connections of the cortex of area 4 of the monkey. Brain: A Journal of Neurology, 101(3), 513–541. 10.1093/brain/101.3.513

Geweke, J. (1982). Measurement of Linear Dependence and Feedback between Multiple Time Series. Journal of the American Statistical Association, 77(378), 304–313. 10.1080/01621459.1982.10477803

Glover, G. H. (2011). Overview of Functional Magnetic Resonance Imaging. Neurosurgery Clinics of North America, 22(2), 133–139. 10.1016/j.nec.2010.11.001

Green, D. M., & Swets, J. A. (1988). Signal Detection Theory and Psychophysics. Peninsula Publishing.

Guo, Z. V., Hart, A. C., & Ramanathan, S. (2009). Optical interrogation of neural circuits in Caenorhabditis elegans. Nature Methods, 6(12), 891–896. 10.1038/nmeth.1397

Histed, M. H., Bonin, V., & Reid, R. C. (2009). Direct Activation of Sparse, Distributed Populations of Cortical Neurons by Electrical Microstimulation. Neuron, 63(4), 508–522. 10.1016/j.neuron.2009.07.016

Hong, N., Vargo, S. M., Hatanaka, G., Gong, Z., Stanis, N., Zhou, J., Belloir, T., Wang, R. K., Bair, W., Chamanzar, M., & Yazdan-Shahmorad, A. (2025). Multimodal Optical Imaging and Modulation through Smart Dura in Non-Human Primates. bioRxiv, 2025.02.27.640384. 10.1101/2025.02.27.640384

Huntley, G. W., & Jones, E. G. (1991). Relationship of intrinsic connections to forelimb movement representations in monkey motor cortex: A correlative anatomic and physiological study. Journal of Neurophysiology, 66(2), 390–413. 10.1152/jn.1991.66.2.390

Izpisua Belmonte, J. C., Callaway, E. M., Caddick, S. J., Churchland, P., Feng, G., Homanics, G. E., Lee, K.-F., Leopold, D. A., Miller, C. T., Mitchell, J. F., Mitalipov, S., Moutri, A. R., Movshon, J. A., Okano, H., Reynolds, J. H., Ringach, D., Sejnowski, T. J., Silva, A. C., Strick, P. L., … Zhang, F. (2015). Brains, genes, and primates. Neuron, 86(3), 617–631. 10.1016/j.neuron.2015.03.021

Kaiju, T., Doi, K., Yokota, M., Watanabe, K., Inoue, M., Ando, H., Takahashi, K., Yoshida, F., Hirata, M., & Suzuki, T. (2017). High Spatiotemporal Resolution ECoG Recording of Somatosensory Evoked Potentials with Flexible Micro-Electrode Arrays. Frontiers in Neural Circuits, 11. 10.3389/fncir.2017.00020

Lepage, K. Q., Ching, S., & Kramer, M. A. (2013). Inferring evoked brain connectivity through adaptive perturbation. Journal of Computational Neuroscience, 34(2), 303–318. 10.1007/s10827-012-0422-8

Li, J., Scholl, L., Le, T., Rajeswaran, P., Orsborn, A., & Shlizerman, E. (2023). AMAG: Additive, Multiplicative and Adaptive Graph Neural Network For Forecasting Neuron Activity. In A. Oh, T. Neumann, A. Globerson, K. Saenko, M. Hardt, & S. Levine (Eds.), Advances in Neural Information Processing Systems (Vol. 36, pp. 8988–9014). Curran Associates, Inc. https://proceedings.neurips.cc/paper_files/paper/2023/file/1c70ba3591d0694a535089e1c25888d7-Paper-Conference.pdf

Liu, Z.-Q., Luppi, A. I., Hansen, J. Y., Tian, Y. E., Zalesky, A., Yeo, B. T. T., Fulcher, B. D., & Misic, B. (2025). Benchmarking methods for mapping functional connectivity in the brain. Nature Methods, 22(7), 1593–1602. 10.1038/s41592-025-02704-4

MacKay, W. A. (2004). Wheels of Motion: Oscillatory Potentials in the Motor Cortex. In Motor Cortex in Voluntary Movements. CRC Press.

Ninomiya, T., Inoue, K., Hoshi, E., & Takada, M. (2019). Layer specificity of inputs from supplementary motor area and dorsal premotor cortex to primary motor cortex in macaque monkeys. Scientific Reports, 9(1), 18230. 10.1038/s41598-019-54220-z

Nolte, G., Bai, O., Wheaton, L., Mari, Z., Vorbach, S., & Hallett, M. (2004). Identifying true brain interaction from EEG data using the imaginary part of coherency. Clinical Neurophysiology, 115(10), 2292–2307. 10.1016/j.clinph.2004.04.029

Nowak, L. G., & Bullier, J. (1998). Axons, but not cell bodies, are activated by electrical stimulation in cortical gray matterI. Evidence from chronaxie measurements. Experimental Brain Research, 118(4), 477–488. 10.1007/s002210050304

Ogawa, S., Fumarola, F., & Mazzucato, L. (2022, May 13). Multi-tasking via baseline control in recurrent neural networks. https://www.semanticscholar.org/paper/74685e98f07d35eced1b709c66a02d2d8fe11257

O’Reilly, J. X., Croxson, P. L., Jbabdi, S., Sallet, J., Noonan, M. P., Mars, R. B., Browning, P. G. F., Wilson, C. R. E., Mitchell, A. S., Miller, K. L., Rushworth, M. F. S., & Baxter, M. G. (2013). Causal effect of disconnection lesions on interhemispheric functional connectivity in rhesus monkeys. Proceedings of the National Academy of Sciences, 110(34), 13982–13987. 10.1073/pnas.1305062110

Orsborn, A. L., Wang, C., Chiang, K., Maharbiz, M. M., Viventi, J., & Pesaran, B. (2015). Semi-chronic chamber system for simultaneous subdural electrocorticography, local field potentials, and spike recordings. 2015 7th International IEEE/EMBS Conference on Neural Engineering (NER), 398–401. 10.1109/NER.2015.7146643

Ouchi, T., & Orsborn, A. L. (2022). Quantifying the influence of stimulation protocols on neural network connectivity inference to optimize rapid network measurements. 2022 44th Annual International Conference of the IEEE Engineering in Medicine & Biology Society (EMBC), 2369–2372. 10.1109/EMBC48229.2022.9871658

Ouchi, T., Scholl, L. R., Rajeswaran, P., Canfield, R. A., Smith, L. I., & Orsborn, A. L. (2025). Mapping eye, arm, and reward information in frontal motor cortices using electrocorticography in non-human primates. The Journal of Neuroscience: The Official Journal of the Society for Neuroscience, e1536242025. 10.1523/JNEUROSCI.1536-24.2025

Oya, H., Howard, M. A., Magnotta, V. A., Kruger, A., Griffiths, T. D., Lemieux, L., Carmichael, D. W., Petkov, C. I., Kawasaki, H., Kovach, C. K., Sutterer, M. J., & Adolphs, R. (2017). Mapping effective connectivity in the human brain with concurrent intracranial electrical stimulation and BOLD-fMRI. Journal of Neuroscience Methods, 277, 101–112. 10.1016/j.jneumeth.2016.12.014

Perich, M. G., Gallego, J. A., & Miller, L. E. (2018). A Neural Population Mechanism for Rapid Learning. Neuron, 100(4), 964–976.e7. 10.1016/j.neuron.2018.09.030

Petkov, C. I., Kikuchi, Y., Milne, A. E., Mishkin, M., Rauschecker, J. P., & Logothetis, N. K. (2015). Different forms of effective connectivity in primate frontotemporal pathways. Nature Communications, 6(1), 6000. 10.1038/ncomms7000

Qiao, S., Sedillo, J. I., Brown, K. A., Ferrentino, B., & Pesaran, B. (2020). A Causal Network Analysis of Neuromodulation in the Mood Processing Network. Neuron, 107(5), 972. 10.1016/j.neuron.2020.06.012

Randi, F., Sharma, A. K., Dvali, S., & Leifer, A. M. (2023). Neural signal propagation atlas of Caenorhabditis elegans. Nature, 623(7986), 406–414. 10.1038/s41586-023-06683-4

Rouse, A. G., Williams, J. J., Wheeler, J. J., & Moran, D. W. (2013). Cortical Adaptation to a Chronic Micro-Electrocorticographic Brain Computer Interface. Journal of Neuroscience, 33(4), 1326–1330. 10.1523/JNEUROSCI.0271-12.2013

Russell, L. E., Dalgleish, H. W. P., Nutbrown, R., Gauld, O. M., Herrmann, D., Fişek, M., Packer, A. M., & Häusser, M. (2022). All-optical interrogation of neural circuits in behaving mice. Nature Protocols, 17(7), 1579–1620. 10.1038/s41596-022-00691-w

Sohal, V. S., Zhang, F., Yizhar, O., & Deisseroth, K. (2009). Parvalbumin neurons and gamma rhythms enhance cortical circuit performance. Nature, 459(7247), 698–702. 10.1038/nature07991

Sun, X., & Xu, W. (2014). Fast Implementation of DeLong’s Algorithm for Comparing the Areas Under Correlated Receiver Operating Characteristic Curves. IEEE Signal Processing Letters, 21(11), 1389–1393. IEEE Signal Processing Letters. 10.1109/LSP.2014.2337313

Sweatt, J. D. (2016). Neural plasticity and behavior – sixty years of conceptual advances. Journal of Neurochemistry, 139(S2), 179–199. 10.1111/jnc.13580

Trumpis, M., Chiang, C.-H., Orsborn, A. L., Bent, B., Li, J., Rogers, J. A., Pesaran, B., Cogan, G., & Viventi, J. (2021). Sufficient sampling for kriging prediction of cortical potential in rat, monkey, and human µECoG. Journal of Neural Engineering, 18(3), 036011. 10.1088/1741-2552/abd460

Valero-Cabré, A., Payne, B. R., Rushmore, J., Lomber, S. G., & Pascual-Leone, A. (2005). Impact of repetitive transcranial magnetic stimulation of the parietal cortex on metabolic brain activity: A 14C-2DG tracing study in the cat. Experimental Brain Research, 163(1), 1–12. 10.1007/s00221-004-2140-6

Wang, X., Gkogkidis, C. A., Iljina, O., Fiederer, L. D. J., Henle, C., Mader, I., Kaminsky, J., Stieglitz, T., Gierthmuehlen, M., & Ball, T. (2017). Mapping the fine structure of cortical activity with different micro-ECoG electrode array geometries. Journal of Neural Engineering, 14(5), 056004. 10.1088/1741-2552/aa785e

Watakabe, A., Ohtsuka, M., Kinoshita, M., Takaji, M., Isa, K., Mizukami, H., Ozawa, K., Isa, T., & Yamamori, T. (2015). Comparative analyses of adeno-associated viral vector serotypes 1, 2, 5, 8 and 9 in marmoset, mouse and macaque cerebral cortex. Neuroscience Research, Marmoset Neuroscience, 93, 144–157. 10.1016/j.neures.2014.09.002

Yang, Y., Connolly, A. T., & Shanechi, M. M. (2018). A control-theoretic system identification framework and a real-time closed-loop clinical simulation testbed for electrical brain stimulation. Journal of Neural Engineering, 15(6), 066007. 10.1088/1741-2552/aad1a8

Yang, Y., Qiao, S., Sani, O. G., Sedillo, J. I., Ferrentino, B., Pesaran, B., & Shanechi, M. M. (2021). Modelling and prediction of the dynamic responses of large-scale brain networks during direct electrical stimulation. Nature Biomedical Engineering, 5(4), 324–345. 10.1038/s41551-020-00666-w

Yazdan-Shahmorad, A., Diaz-Botia, C., Hanson, T. L., Kharazia, V., Ledochowitsch, P., Maharbiz, M. M., & Sabes, P. N. (2016). A Large-Scale Interface for Optogenetic Stimulation and Recording in Nonhuman Primates. Neuron, 89(5), 927–939. 10.1016/j.neuron.2016.01.013

Yazdan-Shahmorad, A., Silversmith, D. B., Kharazia, V., & Sabes, P. N. (2018). Targeted cortical reorganization using optogenetics in non-human primates. eLife, 7, e31034. 10.7554/eLife.31034

Zimmermann, J., Vazquez, Y., Glimcher, P. W., Pesaran, B., & Louie, K. (2016). Oculomatic: High speed, reliable, and accurate open-source eye tracking for humans and non-human primates. Journal of Neuroscience Methods, 270, 138–146. 10.1016/j.jneumeth.2016.06.016

